# New structural insights into the control of the retinoic acid receptors RAR/RXR by DNA, ligands and transcriptional coregulators

**DOI:** 10.1101/2025.03.18.643890

**Authors:** Izabella Tambones, Amin Sagar, Pavla Vankova, Dmitry Loginov, Coralie Carivenc, Natacha Rochel, William Bourguet, Petr Man, Pau Bernadó, Albane le Maire

## Abstract

Retinoic acid receptors (RARs) are ligand-dependent transcription factors essential for various biological processes, including embryogenesis, differentiation, and apoptosis. RARs function as heterodimers with retinoid X receptors (RXRs) and regulate gene expression via retinoic acid response elements (RAREs). Their transcriptional activity is modulated by coregulators, with corepressors maintaining repression in the absence of ligand and coactivators enabling transcription upon ligand binding. Structural studies reveal that DNA binding induces conformational changes affecting coregulator interactions. However, the precise structural organization of RAR/RXR-coregulator complexes and the allosteric influence of DNA on receptor function remain incompletely understood. Our study presents an integrative analysis of the RAR/RXR heterodimer bound to four distinct and relevant RAREs (DR0, DR1, DR5, and IR0) in complex with either a corepressor (NCoR) or a coactivator (TIF-2) nuclear receptor interaction domain. By combining small-angle X-ray scattering, hydrogen/deuterium exchange mass spectrometry and molecular dynamics simulations, we revealed that the heterodimer adopts distinct conformations depending on the DNA sequence, influencing interdomain distances and receptor interactions. Additionally, we uncovered the dynamic interplay between ligand, DNA, and coregulator binding. This study provides new insights into the structural features of coregulator proteins and highlights the allosteric influence of RAREs on receptor function.

## INTRODUCTION

Retinoic acid receptors (RARs or NR1Bs) function as ligand-dependent transcription factors belonging to the nuclear receptors (NRs) superfamily that play multiple central roles in vertebrates, such as embryogenesis, organogenesis, cell growth, differentiation and apoptosis (1). Three RAR subtypes, RARɑ (NR1B1), RARβ (NR1B2) and RARɣ (NR1B3), are encoded by three distinct genes (RARA, RARB and RARG, respectively) (2). Similar to other NRs, RARs exhibit a modular structure with several domains and associated functions, most notably a central DNA-binding domain (DBD), and a C-terminal ligand-binding domain (LBD)(3, 4). RARs form heterodimers with retinoid X receptors (RXRs or NR2Bs) and control the transcription of target genes that contain a retinoic acid response element (RARE) in their promoter region. Naturally occurring RAREs are generally composed of two AGGTCA binding sites arranged in a direct repeat (DR) configuration with a characteristic inter-half-site spacing of one, two or five base pairs (referred to as DR1, DR2 and DR5, respectively) (5–9). Additionally, RAR/RXR can bind a diverse series of RAREs in regulated genes, among them the inverted IR0 sequence and the non-canonical DR0 element that do not act transcriptionally as independent RARE (7). Interestingly, the interaction of RAR/RXR with regions containing the DR0 motif is predominant in undifferentiated pluripotent embryonic cells (8), while in differentiated cells such as MCF-7, the heterodimer-bound regions are characterized by higher frequency of the canonical DR5 sequence (10), highlighting the reorganization of the RAR/RXR binding repertoire during the differentiation process.

Moreover, the biological functions of RARs rely on transcriptional coregulator exchanges mainly controlled by ligand binding (11). In the absence of its cognate ligand, RARɑ (hereafter referred as RAR) represses expression of its target genes by interacting with transcriptional corepressors, like the Nuclear receptor CoRepressor (NCoR/NCOR1) (12), and the Silencing Mediator of Retinoic acid receptor and Thyroid hormone receptor (SMRT/NCOR2) (13) which themselves serve as docking platforms for histone deacetylases that impose a higher order structure on chromatin, not permissive to gene transcription (14). Corepressors interact with NRs through their NR interaction domain (NID), a large disordered region that contains two or three conserved nuclear receptor-binding motifs, so-called CoRNR1, CoRNR2 and CoRNR3 (15, 16). These motifs consist of long amphipathic helices with the consensus sequence LxxxIxx(I/V)Ixxx(Y/F). Among them, CoRNR1 and CoRNR2 serve as the primary motifs of interaction with NRs. In the context of the RAR/RXR heterodimer, RAR and RXR show a preference for CoRNR1 and CoRNR2, respectively (17, 18). The molecular basis of RAR’s repression function has been elucidated through the crystal structure of the RAR-LBD bound to the inverse agonist BMS493 and the CoRNR1 peptide of NCoR (17). The structure reveals that the four-turn helical motif in the NCoR peptide establishes contact within a hydrophobic surface of the RAR LBD formed by the helices H3 and H4, and that the corepressor induces a structural transition of RAR helix H11 into a β-strand conformation (called S3), forming an antiparallel β-sheet with the β-strand of the CoRNR1 N-terminus. Nonetheless, the binding of an agonist ligand - such as all-trans-retinoic acid (ATRA), 9-cis-retinoic acid or the synthetic ligand AM580 – to RAR LBD causes a β-strand (S3) to α-helix (H11) transition, provoking the corepressor release and the repositioning of helix H12, which promotes the recruitment of coactivators such as the steroid receptor coactivator (SRC-1) (19) and the transcriptional intermediary factor 2 (TIF-2) (20). This interaction is established through their NR interaction domain (NRID), characterized by three conserved motifs of LXXLL sequences (21). The coactivator complex formation with the RAR/RXR heterodimer favors the recruitment of histone acetylases and methyltransferases, leading to the chromatin decompaction and target gene transcription.

Over the past 15 years, structural data on integral NRs in various functional states, obtained through a combination of different biophysical and structural methods, have provided a better understanding of the mechanisms involved in molecular regulation. Notably, a limited number of full-length NR structures have unveiled the higher-order organization of these complexes on DNA (LXRβ/RXR (22); PPARγ/RXRα (23); RARβ/RXRα (24); VDR/RXR (25); HNF-4ɑ (26); ERɑ (27); EcR/USP (28); and LRH-1 (29)). Their observed binding modes and conformation revealed relevant interdomain interfaces, proposing that allosteric signals from DNA are passed on to proteins to complete the activation of NRs, and that quaternary structures depend on the receptor type, oligomerization and DNA array array (30–34). In addition, several hydrogen/deuterium exchange mass spectrometry (HDX-MS) studies revealed that DNA binding can induce conformational changes in NR LBD and affect coactivator-NR interactions dynamics, which could play a role in driving promoter specific NR activation profiles (31, 32, 35, 36). Thus, beyond anchoring the NR near the transcription start site of target genes, the architecture of DNA response elements can act as an allosteric regulator of receptor function and its association with coregulators (37). However, the physico-chemical details underlying the assembly and coordination of these large, transient, dynamic macromolecular complexes are yet poorly described.

The low-resolution models of full-length RAR/RXR heterodimers bound to DR5 and DR1 RAREs derived from small-angle X-ray scattering (SAXS) and Förster resonance energy transfer (FRET) revealed that the response elements direct the relative position of the DNA recognition helix and the LBDs, which in turn define the binding site of the coregulators (38). A partial analysis of conformational changes upon DNA binding in the RARβ/RXR heterodimer also revealed regions undergoing different conformational changes upon binding to DNAs with different spacers (DR1 and DR5) that may be linked to transmission through RARβ DBD-LBD interface (24). Interestingly, a recent study has revealed different binding modes and polarity of RAR/RXR heterodimer to DR0 and DR5, leading to differences in conformation and structural dynamics of the RAR/RXR-DNA complexes (39), which could induce differential recruitment of coregulators. These studies suggest that the polarity of the binding motifs and the number of dinucleotide spacers modulate the rotation of the LBDs and the relative positions of the receptors allowed by their highly flexible linkers. However, the precise structural organization of RAR/RXR-coregulator complexes on consensus and non-consensus elements as well as the allosteric influence of specific DNA-binding sites on the recruitment of coactivator or corepressor proteins have not been determined yet. Here, we present a comprehensive analysis combining biophysical, structural, and computational data to investigate the conformations of RAR/RXR heterodimer bound to four relevant DNA response elements (DR0, DR1, DR5, and IR0) and in complex with either the corepressor (NCoRNID) or the coactivator (TIF-2NRID). By combining SAXS data and molecular dynamics (MD) simulations, our integrative approach provides a holistic understanding of the molecular mechanisms and allosteric communication at play in RAR/RXR heterodimer. Our findings reveal that the heterodimer adopts distinct conformations depending on the DNA element, resulting in differential interdomain distances and contacts guided by DNA. The comprehensive analysis of a complete HDX-MS dataset enables us to fully characterize the direct and long-range interactions within the heterodimer and the interconnections among ligand binding, DNA interaction, and coregulator association. Additionally, our analysis reveals new features of NCoRNID and TIF-2NRID in their interaction with RAR/RXR. The insights gained from this extensive study of the complete RAR/RXR complex contributes to a broader understanding of nuclear receptor functions.

## MATERIAL AND METHODS

### Ligands

The ligands AM580, BMS493 and CD3254 were purchased from Tocris Bioscience. All compound stock solutions were prepared at 10 mM in DMSO.

### DNAs preparation

The forward and reverse oligonucleotides corresponding to the different RAREs motifs (DR0, DR0L, DR1, DR5 and IR0) were purchased from Integrated DNA Technologies (IDT) (Supplementary Table 1). The oligonucleotides were dissolved in a TE buffer (20mM Tris pH7.5, 50 mM NaCl) to 1mM concentration, and the complementary strands were mixed in an equimolar ratio. The DNA mix was heated to 95°C for 10-15 minutes and left to cool slowly at room temperature for a few hours. Correct annealing was checked by electrophoretic mobility shift assay (EMSA). For that, ∼100 pmol of DNA was mixed with a gel-loading Dye (New England Biolabs, USA) and loaded onto a 12% acrylamide gel. The run was done in the presence of TBE (1M Tris, 1M Boric Acid, 20mM EDTA) with a 100 V per gel for one hour at 4°C. A similar protocol of double strand DNA preparation was applied for fluorescently-labeled DNA. In this case, the 5’-3’ strand was purchased as Cy5-labeled DNA and mixed with complementary non-labeled DNA in a 5% excess to ensure annealing of the whole Cy5-DNA.

### Constructs, Expression and Purification of Proteins and Complexes

MmRARAΔABF (82-421, Uniprot entry <u>P11416</u>) and HsRXRAΔAB (126-462, Uniprot entry <u>P19793</u>) were cloned into the pET15b and pET28b vectors, respectively, containing an N-ter hexahistidine tag. The recombinant proteins were expressed in the *Escherichia coli* BL21 (DE3) strain. For RAR/RXR heterodimer purification, RAR and RXR were first purified separately. Cells expressing MmRXRAΔAB were harvested and resuspended in binding buffer (50 mM Tris pH7.5, 500 mM NaCl, cOmplete, EDTA-free protease inhibitor cocktail tablet (Roche Applied Science)), sonicated and centrifuged. The supernatant was loaded on a 5 ml HisTrap HP (GE Healthcare). The protein was eluted at 250 mM imidazole and the thrombin tag was cleaved overnight at 4°C after adding 1 unit of thrombin (Sigma Aldrich) per mg of protein. The cleaved protein was further purified by size exclusion chromatography (SEC) on a Superdex 75 26/60 (GE Healthcare) column. The final buffer was 20 mM Tris pH7.5, 150 mM NaCl, 5% (v/v) glycerol, 2 mM DTT. Similarly, cells expressing HsRARAΔABF were harvested and resuspended in binding buffer (20 mM Tris pH8, 500 mM NaCl, 2 mM CHAPS, complemented with EDTA-free protease inhibitor cocktail tablet (Roche Applied Science)), sonicated and centrifuged. The supernatant was loaded on a 5 ml HisTrap HP (GE Healthcare). The protein was eluted at 250 mM imidazole. For the preparation of the RAR/RXR heterodimer, the purified MmRXRAΔAB and HsRARAΔABF were mixed together with a two-fold molar excess of HsRARAΔABF over MmRXRAΔAB and incubated overnight at 4°C. Then, the mix was loaded on a Superdex 200 26/60 (GE Healthcare) column in a buffer consisting in 20mM Tris HCl pH7.5, 75 mM NaCl, 75 mM KCl, 2 mM CHAPS, 4 mM MgCl2, 1 mM TCEP and fractions corresponding to the heterodimer were pooled. Protein samples were concentrated using Amicon-Ultra 30, 000 MWCO centrifugal filter units (Millipore). The purity of the complexes was checked on a SDS-page gel. RAR/RXR heterodimers in complex with DNA response elements were formed by adding a 1.5-fold molar excess of DNA to the receptor dimer followed by an additional SEC step. The NCoRNID (2059-2325, Uniprot entry Q60974) and its mutants were cloned into the pDB-His-TXR-3C vector and expressed using the bacterial BL21(DE3) cellular system. The proteins were first purified using a nickel affinity column in a loading buffer containing 20 mM Tris-HCl (pH 7.5), 300 mM NaCl, and 10 mM imidazole, and eluted with 20 mM Tris-HCl (pH 7.5), 300 mM NaCl, and 500 mM imidazole. This was followed by overnight cleavage at 3°C using 3C protease. The samples were then loaded onto a size-exclusion column (Superdex 75 26/60, GE Healthcare) and purified in 20 mM Tris-HCl (pH 7.5), 150 mM NaCl, and 2 mM dithiothreitol (DTT). The protein-containing fractions were pooled and concentrated using Amicon Ultra-3, 000 MWCO centrifugal filter units (Millipore). The TIF-2NRID (624-773, Uniprot entry Q15596-1) protein was prepared as described by Senicourt and coworkers (40). RAR/RXR heterodimers in complex with NCoRNID or TIF-2NRID were prepared similarly for RAR/RXR LBDs as described by Cordeiro and coworkers (41) and Senicourt and coworkers (40), respectively.

### Steady-State Fluorescence Anisotropy

#### Interaction with DNA in different liganded states

Fluorescent anisotropy assays were performed using a Safire microplate reader (TECAN) with the excitation wavelength set at 635 nm and emission measured at 680 nm for Cy5-labeled DNAs. The buffer solution for assays was 20 mM Tris-HCl, pH 7.5, 75 mM NaCl, 75 mM KCl, 2 mM CHAPS, 4 mM MgCl2, 2 mM TCEP and 10% (v/v) glycerol. We initiated the measurements at the highest protein concentration (2 μM) and diluted the protein sample successively two-fold with the buffer solution. For each point of the titration curve, we mixed the protein sample with 0.5 nM of fluorescent DNA and if stated, with 4 μM of ligands, either AM580 and CD3254 or BMS493 (two molar equivalents). We fitted binding data using a sigmoidal dose-response model (GraphPad Prism, GraphPad Software). The reported data are the average of at least three independent experiments.

#### Interaction of TIF-2NRID with the different DNA-bound heterodimers

Fluorescent anisotropy assays were performed using a Safire microplate reader (TECAN) with the excitation wavelength set at 470 nm and emission measured at 530 nm for Alexa488-labeled TIF-2NRID (40). The buffer solution for assays was 20 mM Tris-HCl, pH 7.5, 75 mM NaCl, 75 mM KCl, 2 mM CHAPS, 4 mM MgCl2, 5 mM DTT and 10% (v/v) glycerol. We initiated the measurements at the highest protein concentration (4 μM) and diluted the protein sample successively two-fold with the buffer solution. For each point of the titration curve, we mixed the protein sample with 10 nM of fluorescent TIF-2NRID and 12 μM of AM580 and CD3254 ligands (three molar equivalents). We fitted binding data using a sigmoidal dose-response model (GraphPad Prism, GraphPad Software). The reported data are the average of at least three independent experiments.

### MicroScale Thermophoresis (MST)

The Atto647N-labeled N-CoR_NID_ was prepared as described in Cordeiro and coworkers (41). RAR/RXR and RAR/RXR-DNA heterodimers were prepared as a twofold serial dilution starting at a concentration of 20 µM in a buffer consisting in 20 mM Tris-HCl, pH 7.5, 75 mM NaCl, 75 mM KCl, 2 mM CHAPS, 4 mM MgCl2, 5% (v/v) glycerol and 0.05 % Tween-20 and added to an equal volume of 80 nM labeled N-CoR_NID_. After 10 min incubation time, the complex was filled into Monolith NT.115 Premium Coated Capillaries (NanoTemper Technologies GmbH) and thermophoresis was measured using a Monolith NT.115 Microscale Thermophoresis device (NanoTemper Technologies GmbH) at an ambient temperature of 22°C, with 5 s/10 s/ 5 s laser off/on/off times, respectively. Instrument parameters were adjusted with 100% red LED power and 40% IR-laser power. Data from three independent measurements were analyzed (NT Analysis software last version, NanoTemper Technologies GmbH) using the signal from Thermophoresis + T-Jump.

### Hydrogen Deuterium Exchange coupled with Mass Spectrometry (HDX-MS)

We used the complexes formed between RAR/RXR heterodimer with different DNAs (DR0, DR1, DR5 and IR0) in the presence and absence of NCoRNID or TIF-2 (Table 1). The HDX reaction was conducted at 4°C where 20µM of the complex sample was diluted 10x into the deuterated buffer (pD 7.5 20mM Tris, 75mM NaCl, 75mM KCl, 4mM MgCl2, 1mM TCEP and 5% glycerol) and stopped after 4*, 10, 90*, 1200 and 10800* seconds (time points denoted with an asterisk were done in duplicate) by adding a quench buffer (0.5M glycine and 4M Urea, pH 2.3) followed by immediate freezing in liquid nitrogen. Samples were stored at -80°C.

**Table 1.** Affinities of DNA and coregulators for the RAR/RXR heterodimer. Dissociation constant (Kd) values for the interaction between the RAR/RXR heterodimer (RR) and the different DNA binding sites were estimated based on fluorescent anisotropy experiments performed in the absence of ligand (apo), in the presence of both the RAR and RXR agonists (AM580+CD3254), or of the RAR inverse agonist (BMS493). Dissociation constant (Kd) values for the interaction between the RR unbound or bound to the different DNA binding sites, wild-type and mutant forms of the corepressor NCoR_NID_ were estimated based on MST experiments performed in the absence of ligand (apo) or in the presence of the inverse agonist (BMS493). Dissociation constant (Kd) values for the interaction between RR unbound or bound to the different DNA binding sites and the coactivator TIF-2_NRID_ were derived from fluorescence anisotropy experiments performed in the presence of the two agonists.

| Protein | DNA | Ligand | K<sub>d</sub> (nM) | Confidence range(nM) |
| --- | --- | --- | --- | --- |
| RR | DR0 | Apo | 37 | 21-66 |
|  |  | AM580+CD325 | 24 | 9-58 |
|  |  | BMS493 | 100 | 45-223 |
|  | DR1 | Apo | 7 | 4-14 |
|  |  | AM580+CD325 | 8 | 5-17 |
|  |  | BMS493 | 27 | 13-56 |
|  | DR5 | Apo | 2 | 1-5 |
|  |  | AM580+CD325 | 2 | 1-4 |
|  |  | BMS493 | 18 | 8-39 |
|  | IR0 | Apo | 3 | 1-6 |

|  |  | AM580+CD325 | 3 | 1-6 |
| --- | --- | --- | --- | --- |
|  |  | BMS493 | 25 | 8-70 |
|  | Complex | Ligand | Kd (nM) | Standard deviation |
| NCOR<sub>NID</sub> | RR | Apo | 490 | 101 |
|  | RR | BMS493 | 368 | 72 |
|  | RR-DR1 | Apo | 531 | 140 |
|  | RR-DR1 | BMS493 | 499 | 124 |
|  | RR-DR1 | AM580+CD3254 | 5300 | 2175 |
|  | RR-DR0 | Apo | 414 | 122 |
|  | RR-DR0 | BMS493 | 244 | 67 |
|  | RR-DR5 | Apo | 283 | 46 |
|  | RR-DR5 | BMS493 | 335 | 123 |
|  | RR-IR0 | Apo | 289 | 76 |
|  | RR-IR0 | BMS493 | 289 | 111 |
| NCOR<sub>NIDmut1</sub> | RR-DR1 | Apo | 347 | 90 |
| NCOR<sub>NIDmut2</sub> | RR-DR5 | Apo | 576 | 155 |
|  | RR-DR5 | BMS493 | 583 | 72 |
| TIF2<sub>NRID</sub> | RR | AM580+CD3254 | 121 | 16 |
|  | RR-DR0 | AM580+CD3254 | 53 | 10 |
|  | RR-DR1 | AM580+CD3254 | 28 | 7 |
|  | RR-DR5 | AM580+CD3254 | 28 | 8 |
|  | RR-IR0 | AM580+CD3254 | 26 | 14 |

Subsequently, each sample was quickly thawed and injected into the LC system with a protease column containing immobilized nepenthesin-2 (42). Digestion was done under a flow rate of 200µL min^−1^ of 0.4% formic acid in the water. The peptides were trapped and desalted on a SecurityGuard pre-column (ULTRA Cartridges UHPLC Fully Porous Polar C18, 2.1 mm, Phenomenex) for 3 minutes. Next, the peptides were separated on an analytical column (LUNA Omega Polar C18 Column, 100 Å, 1.6 µm, 100 mm × 1.0 mm, Phenomenex) using a 6 minutes linear gradient 10–40% of solvent B (A: 2% acetonitrile/0.1% FA in water; B: 98% acetonitrile/0.1% FA in water) (1290 Infinity II LC system, Agilent Technologies). The flow rate on the analytical column was 40 µL min^−1^. The LC system was directly connected to a 15T FT-ICR mass spectrometer’s ESI source (SolariX XR, Bruker Daltonics). The instrument was calibrated externally using sodium TFA clusters and was operated in the broad band mode over the mass range 300-2000 with 1M data acquisition. Data was exported and analyzed using Data Analysis v. 5.3 (Bruker Daltonics) and processed by the custom DeutEx software (43). Data presentation was done using Mstools (44).

Following the separation, the protease column was washed with a 150mM triethyl ammonium acetate, 4M Urea, pH 8.0 solution followed by 5% acetonitrile, 5% isopropanol, and 20% acetic acid solution. Similarly, the trap and analytical columns were washed using 150mM triethyl ammonium acetate/80% acetonitrile, pH 8.0 solution by 1260 Infinity II Quaternary pump (Agilent Technologies).

With the same LC system and gradient elution but an ESI-timsTOF Pro mass spectrometer (Bruker Daltonics, Bremen, Germany), data-dependent LC-MS/MS measurement was used to identify the peptides arising from digestion of each protein. Data were searched using MASCOT v 2.7 (Matrix Science) against a database combining the sequences of RAR, RXR, NCoRNID, TIF-2NRID and contaminants from the cRAP database. Decoy search was enabled and the false discovery ratio was set to 1%. Ion score cut-off was set to 20. Other search parameters were: no-enzyme, precursor mass tolerance 5ppm, fragment ion mass tolerance 0.05Da, no modifications enabled.

### Small-Angle X-ray Scattering (SAXS)

Size-exclusion chromatography coupled small-angle X-ray scattering (SEC-SAXS) experiments were conducted at the SWING beamline on the SOLEIL synchrotron (Saint-Aubin, France) (45). Samples of RAR/RXRΔABF bound to DNA elements DR0, DR0L, DR1, DR5 and IR0, with the ligand BMS493, were analyzed in the absence (RR-DNA complexes) and presence of NCoRNID (NRR-DNA complexes). 50 µL/sample at ∼10 mg/mL was loaded onto a Superdex 200 Increase 10/300 GL column, previously equilibrated with the final buffer (20 mM Tris HCl pH 7.5, 75 mM NaCl, 75 mM KCl, 2 mM CHAPS, 4 mM MgCl₂, 12 mM TCEP), and run at a flow rate of 0.5 mL min^-1^. All experiments were performed at 15°C using CoFlow (at a flow rate of 0.2 mL min^-1^) with the same final buffer to prevent potential aggregation attachment to the capillary. Scattering data were collected with an X-ray wavelength of 1.0332 Å and a sample-to-detector distance of 2 meters. The intensity was measured as a function of the scattering vector, *q*, calculated using the equation: *q* = 4π.sin(θ)/λ, where θ is the scattering angle and λ is the wavelength (46). The collected scattering vectors ranged from *q* = 0.011 to 0.55 Å^-1^. Nine hundred frames were recorded continuously during each run, with a frame duration of 1.99 seconds and a dead time of 0.01 seconds between frames. The 2D scattering images were azimuthally averaged into 1D intensity curves using the Foxtrot software (https://www.synchrotron-soleil.fr/en/beamlines/swing). Homogeneous sample intensity frames were scaled and averaged, and subtraction over the buffer-averaged frames was followed by using CHROMIXS from the ATSAS 3.0.5 software package (47, 48). The final SAXS profiles were used to estimate structural parameters using Primus (49) from the ATSAS suite. The estimated *Rg* via Guinier approximation, *P(r)* and *Dmax* values were calculated using GNOM, and the concentration-independent molecular weight was estimated using Bayesian inference on ATSAS (50, 51).

### Molecular Dynamics Simulation

#### RR-DNA complexes structure preparation

The LBDs for RAR/RXR for all simulations were taken from the crystal structure of the RAR/RXR heterodimer complexed with DR1 (PDB ID: 5UAN). The structures of the DBDs complexed with DNA were obtained from PDB ID: 6XWG for DR5, SASBDB ID: SASDFT8 for DR0, and from the structure solved in this study for IR0. The DNA for DR0L simulations was generated as an ideal B-DNA using the SCFBio web server (https://scfbio-iitd.res.in/software/drugdesign/bdna.jsp). The DBD-LBD linkers were added using the MomaLoopSampler web server (52).

#### NRR-DNA complexes structure preparation

NCoRNID was added to the RAR/RXR-DNA systems prepared above using the structure of the two NCoRNID binding motifs from Cordeiro and coworkers (41). The flexible parts of NCoRNID were then modeled using RanCH and MomaLoopSampler (52, 53). Specifically, RanCH was used to add the flexible terminal segments on the N-terminus of segment 1 and the C-terminus of segment 2 (53). The flexible between the two bound segments was added using MomaLoopSampler(52).

#### Simulation system setup

For all simulations, the hydrogens atoms were added to the DNAs using Chimera before the RAR/RXR-DNA complex was used to setup the simulations using the tools available with Gromacs-2022.6, including addition of hydrogens to the protein atoms to protein residues, solvation and ionization to achieve a final salt concentration of 0.15 M with Na^+^ and Cl^-^. All the simulations were performed using DES-Amber 3.20 force field (54) which has been especially optimized for simulations of DNA-protein complexes. The final prepared system was converted to Amber format using ParmEd (55). At this step, hydrogen mass repartitioning was applied to allow a 4fs time step.

#### Molecular Dynamics Simulations

All simulations were performed using AMBER22 (56). The systems were first minimized for 50, 000 steps, applying position restraints on all protein and DNA atoms. This was followed by NVT equilibration, where the temperature was gradually increased to 303.15 K in 20, 000 steps followed by equilibration at constant temperature for 5 ns. The equilibrated structures were used to run 10 independent NPT simulations, each lasting 1 μs. For each simulation, the first 10 ns were discarded as equilibration.

#### Data analysis

The initial processing of the simulations, including imaging and removal of solvent and ions was performed using CPPTRAJ. Distances and contact maps were calculated usign the MDAnalysis library (57) with custom scripts. For RAR/RXR-DNA simulations, CRYSOL from the ATSAS suite version 3.10 (58) was used to compute the SAXS profile for every 10^th^ structure in the MD trajectories (after discarding the first 10 ns). The resulting SAXS profiles were reweighted using Bayesian Optimization approach, as implemented in BioEn (59). The weights obtained from Bayesian reweighting were used to generate both reweighted distance distributions and reweighted contact maps.

## RESULTS

### DNA binding affinity to the RAR/RXR heterodimer in active and repressive states

To assess and compare the RAR/RXR heterodimer’s binding affinities to distinct RARE binding sites, we performed fluorescence anisotropy measurements in the presence and absence of the cognate ligands. For this, we expressed and purified the RARαΔABF-RXRαΔAB heterodimer (referred to as RAR/RXR or RR when in complex) without the NR flexible regions (the N-terminal A/B and C-terminal F domains) (Figure S1A). Biophysical characterization by SEC-MALS ensured the sample integrity (Figure S1B). The heterodimer was then incubated with either both agonists (AM580 for RAR and CD3254 for RXR) or the RAR inverse agonist ligand (BMS493), to reinforce either the active or the repressive conformational states, respectively. We used four different canonical and non-canonical CY5-labeled RAREs binding sites (DR0, DR1, DR5 and IR0 - Table S1), chosen according to their relevance and binding prevalence for the RAR/RXR heterodimer (7, 9, 24, 39). The measured affinities were in the nanomolar range (Table 1, Figure S2A), confirming the high binding affinity of RAR/RXR with all tested DNA sequences (7, 8, 24, 60). Unliganded RAR/RXR exhibited the highest affinity for IR0 (Kd = 3 nM) and DR5 (Kd = 2 nM), followed by DR1 (Kd = 7 nM), with a significantly lower affinity for DR0 (Kd = 37 nM). These results were consistent with previous observations for similar DNA sequences (7, 39). While the affinity pattern was maintained under unliganded and agonist-bound conditions, affinities measured in the presence of BMS493 were slightly reduced, likely due to the destabilizing effect of the inverse agonist ligand on the heterodimer.

### DNA-Driven Structural Adaptations of the RAR/RXR Heterodimer

Seeking to explore how the DNA binding could influence the conformational states of the RAR/RXR heterodimer, we conducted SEC-SAXS experiments on the various RR-DNA complexes. The scattering intensities as a function of the momentum transfer vector, *Q*, for all conditions are displayed in Figure 1A-D. No deviations from linearity were observed in the Guinier region of all DNA conditions, indicating no sample aggregation or interparticle effects (Figure 1A–D, inset). All the studied complexes presented a radius of gyration (*R_g_*) of approximately 4.0 ± 0.4 nm, as determined by the Guinier approximation, and the pairwise distance distribution functions, *P(r)*, indicated a maximum particle dimension (*D_max_*) around 13.0 ± 0.9 nm, regardless of the DNA element type (Table S2). Despite SAXS-derived parameters pointing to no significant variations of the overall dimensions related to the type of bound DNA, a disparity in *P(r)* and Kratky profiles among the RR-DNA complexes was observed (Figures 1E–H and S3A-F). All *P(r)* curves displayed distinct asymmetric patterns, showing a maximum at 3 nm followed by a hump that extended into an elongated tail at higher r values (5–8 nm), indicating the coexistence of compact and extended RAR/RXR conformations that varied according to the DNA type. For RR-IR0, a prominent peak at 3 nm was followed by a second peak at 7 nm, consistent with the *P(r)* function of a dumbbell-shaped particle (61), indicating two well-separated centers of mass corresponding to the more distantly positioned RAR/RXR DBDs and LBDs (Figure 1H). RR-DR5 and RR-DR1 complexes shared a similar tendency, even if less pronounced for the second peak, suggesting less defined conformation (Figure 1F-G). In contrast, the RR-DR0 profile exhibited a single peak, confirming a predominant compact state (Figure 1E, gray), in line with previous reports (39). Given RR-DR0’s contrasting behavior, we conducted molecular dynamics (MD) simulations on this complex to further characterize its structural properties (detailed below). Our simulations suggested that the heterodimer tended to adopt a highly compact but non-physical conformation when bound to DR0, likely due to the short length of this DNA (16 nucleotides) combined with its topology. To ensure relevant states, we next used a longer DR0 sequence with 30 nucleotides (hereafter referred to as DR0L) for the SAXS analysis (Figure 1A, light red). The structural parameters obtained for RR-DR0L were in a similar range as observed for other DNA element conditions (Table S2). Additionally, its *P(r)* presented a broader peak than previously observed for DR0, with distances ranging from 3-6 nm, yet assuming its particular dynamic behavior and contrasts with the other DNA elements. Overall, the SAXS profiles of the RR-DNA complexes suggested that the RAR/RXR heterodimer displays distinct structural and dynamic signatures depending on the bound DNA motif.

**Figure 1.**
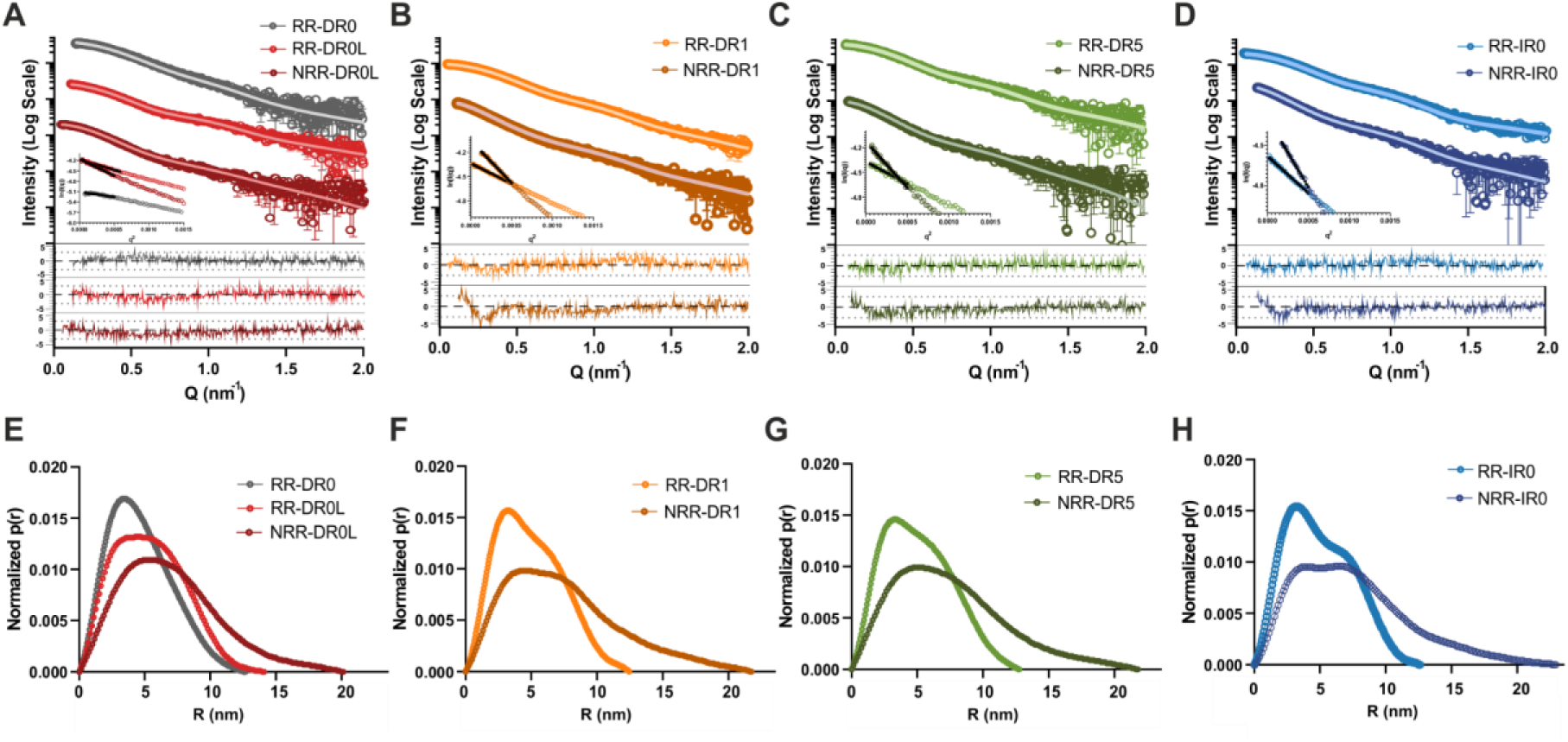
SEC-SAXS data measured on RAR/RXR Complexes Bound to Different DNA Binding Sites in the Presence and Absence of NCoRNID. **(A-D)** SAXS intensity profiles for RAR/RXR complexes bound to different DNA binding sites (RR-DNA) (Panel A: DR0 and DR0L, Panel B: DR1, Panel C: DR5 and Panel D: IR0). Color-coded lines indicate each condition in the absence (RR-DNAs, light colors) and presence of NCoRNID (NRR-DNAs, dark colors). Insets illustrate the Guinier plot for each condition. Below, the normalized residuals are displayed, estimated from the SAXS fitting. **(E-H)** Pairwise distance distribution functions *P(r)* derived from SAXS data, using the same color pattern as in panels A-D for RR-DNA and NRR-DNA complexes. The *P(r)* distribution are shown for heterodimers bound to RR-DR0, RR-DR0L and NRR-DR0L (Panel E); RR-DR1 and NRR-DR1 (Panel F); RR-DR5 and NRR-DR5 (Panel G); RR-IR0 and NRR-IR0 (Panel H).

To further describe the conformations and dynamics of RR-DNA complexes, we conducted all-atom MD simulations running 10 independent 1 µs trajectories per DNA condition using the DES-Amber-3.20 force field, which was developed to sample the conformational landscapes of protein-DNA complexes (62). For that, the heterodimer was assembled on each DNA, accounting for already described polarization and specific binding modes for RAR/RXR bound to DR1, DR5 and DR0 (24, 39). To build the model of RAR/RXR bound to IR0, we attempted to crystallize the RAR/RXR DBD heterodimer assembled on IR0, but only crystals of the IR0-bound RXR DBD homodimer were obtained. The RXR/RXR IR0 crystallized in the P4_1_2_1_2 space group, with one dimer-DNA complex per asymmetric unit. The two RXR DBDs were bound to opposite sides of the DNA, and no interaction is observed between them (Figure S4-A), consistent with the binding to IR0 as two monomers at equivalent sites in a head-to-head fashion with the T-boxes of each monomer pointing towards the 3’- and 5’-flanking sequences of the IR0. Each half-site made specific contacts with 4 base pairs (Figure S4-B and C). Both RXR DBDs also formed contacts with the phosphate backbone. To determine the polarity of the DBDs bound to IR0, we performed FRET experiments using a TAMRA5/6-labeled DNA and an acceptor-labeled polyHis-N-ter-RAR with OG488-TriNTA and purified RARΔAB/RXRΔAB. The value of the FRET efficiency at saturation (Esat ∼0.7) (Table S3) that was similar to the values obtained for Ramp2 DR1 and Hoxb13 DR0 (39), suggesting a binding mode that favored a close proximity between the donor and acceptor dyes and thus placed RAR bound to the 5’ half site (Figure S4-D).

Across all the RR-DNA simulations, we observed that the folded regions od RAR/RXR remained well-ordered and formed a stable complex with the DNA. Next, we computed the SAXS theoretical intensity profiles for the four RR-DNA complex trajectories. The excellent fitting between the averaged theoretical profiles and the experimental SAXS data for the RR-DR5 complex highlighted the robustness of our simulation model for this DNA condition (Figure 1D, Table S2). An additional step of Bayesian reweighting (59) was required to further optimize the fitting obtained for the RR-DR0L, RR-DR1 and RR-IR0 complexes (Figure 1A, B and D, Table S2). Building on the excellent agreement between experimental and simulated data, with χ² values ranging from 1.0 to 1.4, we conducted an in-depth analysis of interdomain distances and intra- and inter-molecular contacts within the simulated DNA-bound heterodimers. The distances measured between the DBD and LBD of RAR/RXR bound to the different DNA elements ranged from 4 to 8 nm (Figure S3G). Moreover, by looking into the individual domains in the complexes, we observed that in RR-DR0L and RR-DR1, RAR adopted preferentially shorter DBD-LBD distances, with a maximum density of ≈6 nm (Figure 2A-B, left). In parallel, the high frequency of short distances between RAR DBD and RXR LBD highlighted the compact configuration across subunits imposed by RAR within these complexes (Figure 2A-B, right). Conversely, in RR-DR5, RAR shifted to an elongated yet heterogeneous conformational behavior, and in RR-IR0, RAR shifted the conformational equilibrium to high distances whereas RXR domains were preferentially closely positioned (Figure 2C-D). Overall, our results indicate that each DNA differently drives the relative subunit domain positioning of RAR/RXR heterodimer.

**Figure 2.**
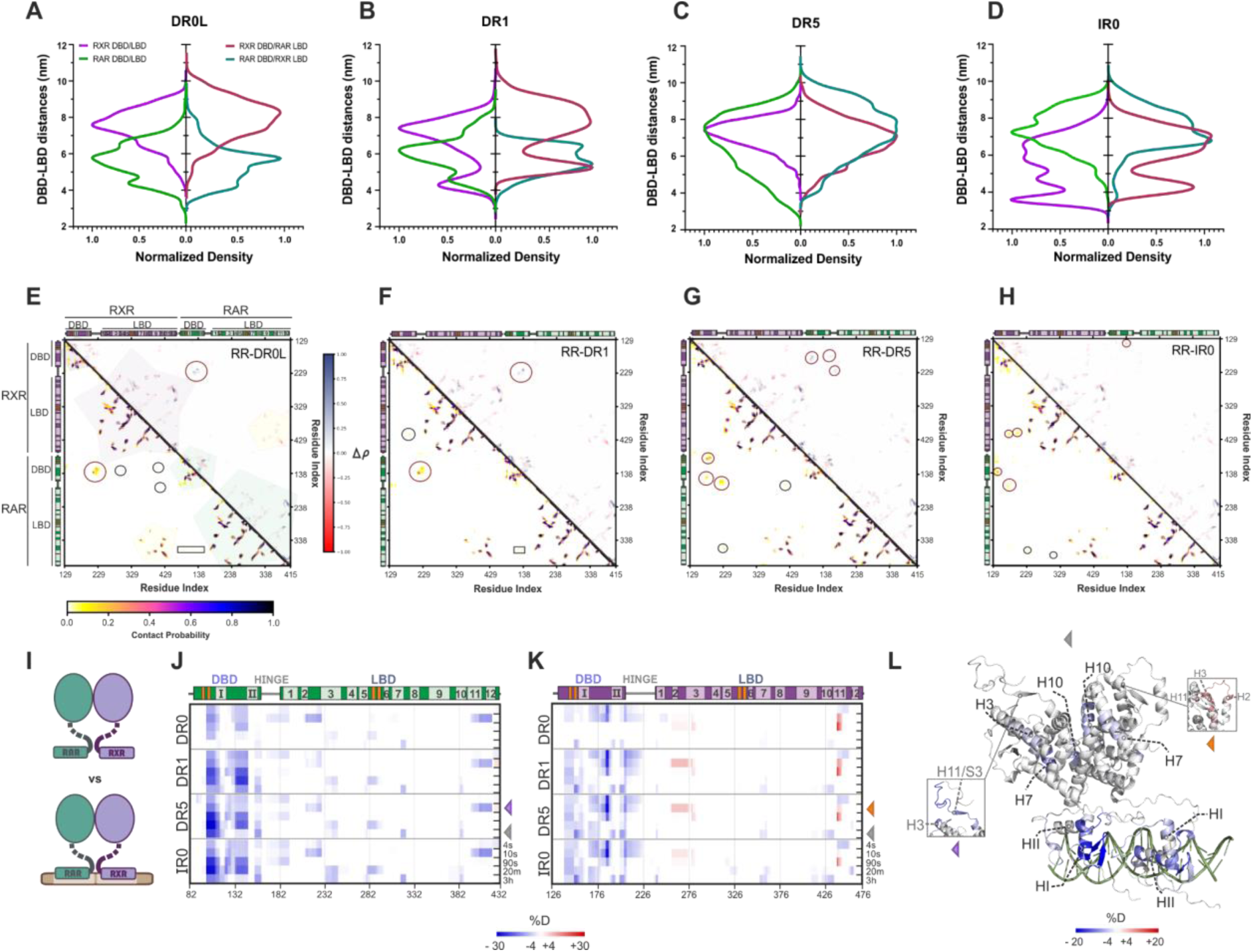
Molecular Dynamics and H/D Exchange based Analysis of RAR/RXR Complexes Bound to Different DNA Elements. **(A-D)** Interdomain distances between DBD and LBD within RAR/RXR complexes derived from reweighted simulations. The left-side graph in each panel shows the interdomain distances for RAR (purple) and RXR (green) receptors, whereas the right-side graph displays distances between RXR DBD and RAR LBD (dark pink) and RAR DBD and RXR LBD (dark green). Panels are organized by DNA binding site: DR0L (Panel A), DR1 (Panel B), DR5 (Panel C), and IR0 (Panel D). **(E-H)** Lower-left: Contact probability maps for RR-DNAs complexes generated from the reweighted simulations. Up-right: Differential plots of normalized weighted contact densities (Δρ) calculated as Δρ = ρRR-DNA − ρRR. Positive Δρ values (blue) represent regions where the contact density increases in the presence of DNA, and negative (red) represent regions where it decreases. Protein secondary structures are indicated in colored bars: RXR (purple) and RAR (green), with alpha-helices (light shades) and beta-sheets (orange). Polygons highlight the intradomain contacts (purple for RXR and green for RAR), with yellow polygons indicating LBD heterodimerization contacts. Circles highlight the specific contacts that vary according to DNA conditions, while rectangles indicate DBD-LBD contacts within each receptor. Panels correspond to DNA binding sites (Panel E: DR0L, Panel F: DR1, Panel G: DR5, and Panel H: IR0). **(I)** Illustration comparing the apo RAR/RXR state vs. apo RAR/RXR bound to DNA. **(J)** Differential heatmap highlighting structural changes in RAR and **(K)** RXR in the RAR/RXR heterodimer upon binding of different DNAs. Deuteration level of the apo forms of either RAR or RXR were subtracted from those obtained for both proteins in the presence of the DNA ligands. Each thick bar represents one experimental condition and can be subdivided into thinner bars representing the time points of exchange (indicated on the right). Structural elements are shown above the plots. **(L)** Representative H/D exchange data corresponding to the binding of RAR/RXR heterodimer to DR5 plotted on the model generated by AlphaFold 3.0 (63). The data corresponds to exchange times of 3 hours, highlighted on the right side of the heatmaps by the gray arrowhead. The inset shows additional dynamics (exchange time 10s) of DNA-induced protection of RAR regions involved in coregulator interfaces (purple arrowhead) and the antagonist effect on RXR, leading to deprotection at its coregulator interface (orange arrowhead).

Since domain positioning can influence interdomain interfaces, we estimated residue contact probabilities within the RR-DNAs complexes (Figure 2E-H). As expected, all complexes demonstrated strong intradomain contacts as well as in the LBDs’ heterodimerization surface. The heterodimers bound to DR0L and DR1 particularly presented specific contact probabilities involving the RAR DBD with the RXR hinge and, to a lesser extent, with both LBDs (Figure 2E-F). The contact submatrix generated for residues previously identified as part of the RAR DBD-LBD interface (HI-HII loop and H9-H10 loop, respectively) in studies of full-length NRs (24, 64, 65) revealed the prevalence of interaction within RAR bound to DR0L and DR1 binding sites, albeit transiently (Figure S3H-K). These findings aligned with the model proposed by Chandra and colleagues (24), where the RAR DBD and LBD contacts allowed allosteric signal transmission, and were essential for the receptor’s repressive response on DR1 elements. In contrast, in RR-DR5 and RR-IR0, RAR lacked the DBD-LBD interface, in line with increased interdomain distances (Figures 2G-H and S3J-K). Residues in the RAR LBD (H9-H10 loop) participated in alternative interfaces, including interactions with the RXR hinge. In contrast, RXR retained sufficient compactness in RR-IR0, facilitating interdomain communication and showing contact between its LBD H9/H10 residues and the HI-HII region of its DBD (Figure 2H). Furthermore, we observed variations in hinge-mediated contacts with the rest of the heterodimer. In RR-DR1 and RR-DR0, the RAR hinge interacted with RAR DBD and LBD; however, these contacts were abrogated in other complexes. Moreover, the RXR hinge exhibited clear interactions with RAR DBD in RR-DR0 and RR-DR1 but shifted to a hinge-hinge interaction in RR-DR5 and RR-IR0. To note, these variations appeared to arise from specific DNA binding (Figure 2E-H, upper-right). In RR-IR0 complexes, the compact conformation of RXR allowed its DBD to establish transient interactions with the RAR hinge. In conclusion, our models showed that compact complexes centered around RAR (e.g., RR-DR0 and RR-DR1) exhibited a higher probability of long-range interdomain contacts within this receptor. In contrast, extended complexes lost these DBD-LBD interactions, preferentially retaining contacts between neighboring regions (for the RR-DR5 complex) or within RXR (for RR-IR0). Our findings support a model in which the RAR/RXR heterodimer adapts its conformation to specific DNA binding sites by modulating domain positioning and shaping its contact repertoire, potentially influencing coregulator interactions.

### Long-Range Allosteric Effects in the RAR/RXR Heterodimer Upon DNA Binding

We then experimentally investigated the direct and long-range effects induced by DNAs on the RAR/RXR heterodimer. To this end, we employed HDX-MS to monitor dynamic conformational changes in the different RAR and RXR domains upon DNA binding. This approach tracked altered backbone amide hydrogen exchange with deuterated solvent across different time points, enabling the detection of short- and long-term exchanges. The RR-DNA complexes were prepared without ligands to better approximate native conditions and accurately reflect the cellular context (Figure S1C). After obtaining a nearly complete sequence coverage of RAR and RXR (Figures S5 and S6), we calculated the differences in HDX induced by the binding to the different DNA elements (Figure 2I). Our results showed that direct contact with the DNA promoted protection of RAR and RXR DBDs (Figure 2J, K). More specifically, we observed strong protection on the β1/β2, the N-terminus of α-helix HI and the loop HI-HII of the DBDs, consistent with their regions of insertion into the DNA. No protected surface differences were detected at the DBD heterodimerization regions despite structural models and contact maps indicating variations on the HI-HII loop and hinge, depending on the bound DNA element (Figure 2E-H). The overlap between DNA-induced heterodimerization residues in the DBDs and those involved in nucleic acid interactions (Figure S3L) likely resulted in mixed protections, making it challenging to distinguish specific variations in this region using HDX-MS. Nevertheless, differences in protection intensities in the DBDs were observed among the DNA elements. Specifically, the RAR and RXR DBDs bound to DR0 exhibited a less pronounced protective effect, restricted to shorter time points, suggesting a more transient and weaker DNA interaction than for other complexes (Figures 2J-K and S7A). This observation aligned with the lower binding affinity measured for DR0 (Table 1). Furthermore, upon all DNA binding, the RXR DBD exhibited a broader protection area than the RAR DBD, consistent with the larger interaction surface observed in the crystallographic structure (66). RXR showed diffuse and less intense protection, suggesting that RAR was the primary anchor point of the heterodimer on DNA. This observation aligned with findings on other RXR partners, such as VDR and FXR (36, 67).

Although less pronounced than in the DBD, the DNA binding also influenced more distant regions of the heterodimer, including the hinge and LBDs (Figure 2J-K). Upon DNA binding, the RAR hinge exhibited short-term protection, ranging from 4 to 90 seconds, in agreement with the hinge-mediated contacts previously observed in MD simulations. Furthermore, DNA induced significant protection in a defined region of the RAR LBD that included the hinge C-terminus (aa 162–197), H3 (aa 217–242), the H5/β-sheet segment (aa 267–292), and H11 (aa 387–412), which are regions known to participate in ligand binding and coregulator interactions (Figure 2L, inset). Further protection was also observed on RAR LBD heterodimerization interface, especially on H7 (aa 307-317) and H10 (aa 372-382) (Figure 2J). These results matched the similar protective effect observed on RXR LBD’s H7 (aa 348-363) and H10 (aa 408-428) (Figure 2K), in agreement with a stabilization of the heterodimerization interface when RAR/RXR was bound to DNA (Figure 2L). Interestingly, we observed that DNA binding likely caused a substantial destabilization to RXR LBD’s H2-H3 loop (aa 253-275) and H11 (aa 428-443), corresponding to the surface of interaction with coregulators (Figure 2K-L, inset). These results suggest an antagonistic role of DNA on the heterodimer subunit LBDs and reinforce RXR’s subordinate role. Altogether, our findings highlight that DNA influences RAR/RXR domains through direct interactions and long-range allosteric effects. These effects may affect the heterodimer functions, including heterodimerization stability, ligand binding and coregulator interaction.

### DNA Conformational Signature Persists During Repressive Complex Formation with NCoRNID

In the absence of its cognate ligand, RAR/RXR bound to RAREs formed a repressive complex by recruiting corepressor molecules such as NCoR. We tested if and how DNA could play a role in the interactions within these repressive complexes. Thus, to quantify the interaction affinity between the different RR-DNA complexes and the corepressor NCoRNID, we performed microscale thermophoresis (MST) experiments, in the presence and absence of the BMS493 ligand. Our measurements indicated an affinity of NCoRNID for RAR/RXR around Kd = 500 nM, with the interaction enhanced in the presence of BMS493 (Table 1, Fig S2B). In contrast, the affinity was drastically reduced in the presence of the RAR agonist ligand (AM580) with a Kd superior to 5 µM, as expected. These affinities were in agreement with the previous measurement conducted by MST on RAR/RXR LBDs (41). Interestingly, the NCoRNID affinity was similar for unliganded RR-DR0 and RR-DR1 complexes, whereas slightly enhanced for the RR-DR5 and RR-IR0 (Table 1, Figure S2C). An increase in affinity was observed in the presence of BMS493, except for RR-DR5 and RR-IR0. Our results suggested that DNA binding induces variations in corepressor affinity with RAR/RXR and in responsiveness to ligands.

To explore the structural and dynamic basis of these variations, we first investigated whether NCoRNID binding influences the DNA-dependent conformational patterns of the RAR/RXR heterodimer. For this purpose, we collected SEC-SAXS data on the RAR/RXR heterodimer bound to NCoRNID in the presence of different DNA elements (NRR-DNA complexes). After the addition of NCoRNID, the Rg of the complexes increased by 1.5 nm (to approximately 5.5 ± 0.2 nm), and the *Dmax* expanded to 21.6 ± 1.5 nm (Table S2), consistent with the NCoRNID contribution previously observed in RAR/RXR LBDs (41). Similarly, the *P(r)* profiles for the NRR-DNA complexes showed that NCoRNID shifted pairwise distance frequencies toward higher r values across all conditions, reflecting the enlargement of the overall assembly size (Figure 1E–H). Furthermore, adding NCoRNID increased the system flexibility, reducing the resolution between shorter and longer distance peaks. Despite this, the superimposed SAXS-derived plots (*P(r)* and Kratky profiles) maintained the variations observed across DNA conditions, consistent with the DNA-specific signatures previously described for the RR-DNA complexes (Figure S3M-N).

Subsequently, we performed 10 independent 0.5 µs all-atom MD simulations of RR-DNA in complex with NCoRNID binding to both the RAR and RXR (deck model) and generated models of NRR-DNA complexes by using simulated heterodimer, followed by the addition of NCoRNID bound to RAR (asymmetric model), previously generated by Cordeiro and coworkers (41). We re-weighted and fitted the SAXS theoretical curves from the trajectories and models with the experimental intensity curves (Figure 1A-D). Here again, the excellent fitting, with χ² values ranging from 1.3 to 2.1, enabled the in-depth analysis of the reweighted simulations of these complexes. Firstly, we estimated the DBD-LBD distances within the NRR-DNA complexes and calculated the differential variations relative to the RR-DNA complexes (Figure 3A-D). Our results showed that under repressive complex formation, RXR preferentially shifted its interdomain frequency toward longer distances. This effect was pronounced in RR-DR1 and RR-IR0 complexes and was also observed in crossed distances (RXR DBD/RAR LBD, Figure 3B and D). The minimal impact of the corepressor interaction on RAR suggests that RXR was primarily responsible for undergoing the conformational adjustments required to sustain optimal interaction with NCoRNID.

**Figure 3.**
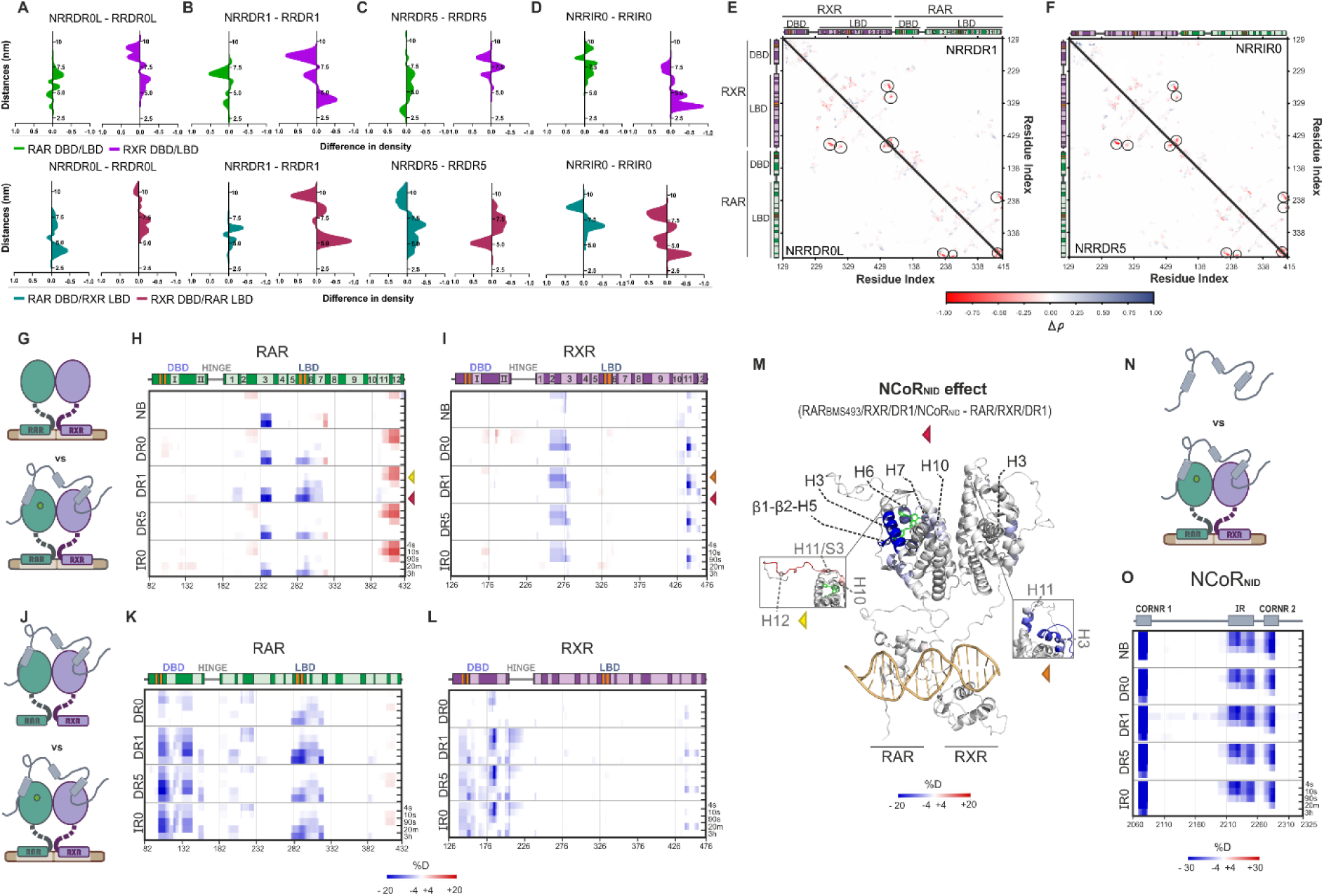
The interaction of RAR-BMS493/RXR bound to DNA with the NCoRNID reveals new features. **(A-D)** The differential on interdomain distances induced in RAR/RXR upon NCoRNID interaction (NRRDNAs - RRDNA). Light green indicates differences in distances for RAR DBD-LBD, purple represents RXR, dark green corresponds to RAR-DBD/RXR-LBD, and red denotes RXR-DBD/RAR-LBD. Panels are organized by the bound element (Panel A: DR0L, Panel B: DR1, Panel C: DR5, Panel D: IR0). **(E-F)** Differential plots of normalized weighted contact densities (Δρ) calculated as Δρ = ρNRR-DNA − ρRRDNA. Positive Δρ values (blue) represent regions where the contact density increases in the presence of NCoRNID, while negative (red) indicate regions where it decreases. (E) The lower-left quadrant corresponds to NRR-DR0L, while the upper-right quadrant represents NRR-DR1. (F) The lower-left quadrant corresponds to NRR-DR5, and the upper-right quadrant represents NRR-IR0. **(G)** Illustration comparing apo RAR/RXR-DNA vs RAR-BMS493/RXR/DNA bound to NCoRNID. RAR is depicted in green, RXR is shown in purple, and NCoRNID is represented in gray. The RAR inverse agonist (BMS493) is drawn as a light green circle, and the DNA is shown as brown bars. **(H)** Differential heatmap highlighting structural changes in RAR (apo - NB and DNA bound forms) upon BMS493 and NCoRNID binding. Red and blue shades indicate increased and decreased deuterium incorporation, respectively. Secondary structural elements are shown above each heat map, where α-helices are represented in lighter colors and labeled with their respective numbering, and β-strands are shown in orange. **(I)** Same structural effects as in (H) but on the RXR part of the RAR/RXR heterodimer. **(J)** Illustration comparing RAR/RXR-NCoR with RAR-BMS493/RXR-NCoRNID bound to the DNAs. **(K-L)** Differential heatmaps showing structural changes in RAR and RXR, respectively, based on the conditions illustrated in (J). **(M)** HDX differential protections plotted on the model generated from the crystallographic structure of RARβ/RXR-DR1 (24). The data corresponds to exchange times of 3 hours and 10 seconds (inset) - highlighted on the right side of the heatmaps by colored arrowheads. **(N)** Illustration of the NCoRNID vs the NCoRNID bound to RR-DNAs. **(O)** Differential HDX-MS heatmap of NCoRNID, showing the impact of the RAR-BMS493/RXR/DNA interaction on the corepressor protection.

To identify contact variations associated with corepressor binding, we calculated the differential contact probabilities within the RAR/RXR heterodimer in NRR-DNA relative to RR-DNA complexes (Figure 3E-F). In all conditions, NCoRNID abrogated specific intradomain contact probabilities on the RXR and RAR C-terminus (including H12), H3 and H4 of their LBD. This loss of intramolecular contacts is in agreement with the structural changes in RAR and RXR required to form the interface with the corepressor. These results aligned with the expected binding pattern of the corepressor, where CoRNR1 interacted with the RAR LBD (H3-H4 and H11), while CoRNR2 interacted with the RXR LBD (H3-H4 and H12) (Figure S3O-R) (17, 41). Beyond these variations, no significant changes in contact probabilities were observed in the NRR-DNA complexes. Notably, the LBD-DBD contacts observed for RAR bound to DR0L and DR1 persisted in the presence of the corepressor (Figure S3S-V). Overall, our findings capture the primary interface and dynamic effects driven by NCoRNID and confirm that the conformational variations in the RAR/RXR heterodimer induced by DNA binding remain in the repressive complex. Despite that, additional RXR conformational changes may still occur to optimize interactions with the corepressor.

### Structural and Functional Characterization of NCoRNID Interactions with RAR/RXR bound to different DNA Elements

To track the direct and long-range effects induced by NCoRNID and DNA binding and gain insight into their interplay, we performed HDX-MS experiments on RR and RR-DNA complexes in the presence of NCoRNID. Due to the disordered nature of NCoRNID transient interactions, it was crucial to run the experiments at low temperature (4°C) to slow down the exchange and collect very short exchange times. Details into the sequence coverage for the NRR-DNA complex are available as supplementary material (Figures S5, S6 and S8). The addition of BMS493 to the NRR-DNA reinforced the repressive interactions, avoiding the complex dissociation during sample preparation and data collection. Additionally, the presence of the ligand allowed the visualization of the DNA impact on ligand binding under repressive conditions. Prior to HDX-MS experiments, the integrity of the complex was checked by SDS-PAGE and SEC-MALS (Figure S1B and D).

We first report the structural changes of H/D exchange in RR and RR-DNA complexes induced by the interaction with the ligand and NCoRNID (Figure 3G-I). NCoRNID binding did not significantly affect the heterodimer DBDs (Figure 3H-I). However, it impacted H/D exchange protection in both RAR and RXR LBDs through their well described interaction interface. Notably, adding the corepressor induced strong protection on RAR H3, for all RR-DNA complexes (Figure 3H and M). As part of the classical interaction surface with the NCoRNID, RAR H11 and H12 showed a significant deprotection in the presence of NCoRNID (Figure 3H and M, yellow arrowhead). This deprotection occurred due to the differences in deuterium incorporation of the H11 and the β-strand (S3) registering the structural secondary transition in response to NCoRNID binding, as previously described by X-ray crystallography (17) and in agreement with the contact probabilities observed in MD trajectories of the NRR-DNA complexes (Figure S3O-R). Of note, this deprotection was also connected to the increased H12 dynamics as part of the repressive signature on RAR, and it is further enhanced in the presence of DNA and ligand. The RAR inverse agonist ligand (BMS493) presented clear protective effects in the RAR ligand binding pocket (LBP; β1/β2-H7) (Figure 3H and K), notably with stronger protection for NRR-DR1.

Differences in H/D exchange protection in RR-NCoRNID following DNA binding (Figure 3J-L) revealed protective signals originating in the DBDs and extending through the RAR LBD H1 to the β1/β2-H7 region, with pronounced protection observed for the DR1 complex (Figure 3K). These results align with the protection observed in RR-DNA complexes without NCoRNID (Figure 2J), indicating that the DNA effects on RAR were maintained upon interaction with the ligand and corepressor, contributing to a stable complex formation. On RXR side, significant protections were observed in its LBD following NCoRNID binding (Figure 3I). Specifically, corepressor binding induced protection in the H2-H3 and H11-H12 regions of the RXR LBD, indicating that NCoRNID interacted with both receptors within the heterodimer (deck model). Finally, the DNA destabilization played on RXR LBD previously observed for the RR-DNA complexes (Figure 2K) appeared abrogated in the presence of NCoRNID (Figure 3L), suggesting also an overall positive stabilization of NCoRNID on RXR LBD. The differences in protection between the receptors confirmed RAR as the primary interaction interface with NCoRNID and a synergistic stabilizing effect of DNA and NCoRNID. Relative to other elements, DR1 displayed the strongest protective effect on the corepressor and ligand interface (Figure S7B), highlighting the specific role of this DNA in stabilizing the RAR/RXR repressive complex through long-range effects. These findings could explain the repressiveness response of RAR/RXR complexes on DR1 elements, as previously observed (24).

Subsequently, we analyzed the effects of RR and RR-DNA complexes on NCoRNID to identify its interacting regions (Figure 3N). In the presence of RR, the corepressor exhibited the strongest protection on CoRNR1 (Figure 3O), which serves as the primary anchor point through direct binding with RAR. Additionally, this protection was enhanced in the presence of the ligand and DNA (Figure S7C). Interestingly, DR1 provided the strongest protection for CoRNR1, demonstrating DNA’s ability to affect coregulator stability when bound to RAR/RXR. In contrast, the additional significant protection observed in the CoRNR2 region of NCoRNID did not intensify upon DNA binding. As previously reported, this motif may interact with RXR, supporting NCoRNID binding to RAR/RXR through its two motifs (41). In this model, CoRNR2 formed a weaker, likely transient interaction interface with RXR, further destabilized by the presence of DNA. Unexpectedly, the RR binding generated explicit protection on the intermediate region (IR) of NCoRNID (Figure 3O), which was not significantly modulated by DNA nor by the ligand (Figure S7C). This IR region (aa 2209–2254), located between CoRNR1 and CoRNR2, was predicted to adopt a helical structure and was highly conserved among corepressors. Furthermore, it was previously proposed to primarily mediate intramolecular interactions (41).

To gain deeper insight into the role of IR in the overall interaction of NCoRNID with the RAR/RXR heterodimer, we modeled the repressive complex using AlphaFold3.0 (63). The NRR-DR1 model, featuring an IR region composed of two α-helices, was subjected to 0.5 µs all-atom MD simulations (10 independent runs) to assess its stability over time and potential gain of interactions with RAR or RXR. Our simulations revealed the stable α-helix secondary state of the IR region across the simulations (Figure S9A). Furthermore, enhanced interactions with IR of NCoRNID primarily involved RXR, mainly through its H11/H12 region neighboring the CoRNR2 interface of interaction (Figure S9B). This new interface of interaction appeared to have reduced local interaction time compared to CoRNR1 and CoRNR2 with the heterodimer (Figure S9C), corroborating the H/D differential in protection intensities (Figure 3O). Next, to assess the impact of the IR region on the interaction between NCoRNID and the heterodimer, we produced two NCoRNID mutants: one in which the IR region was replaced by a fully disordered segment (NCoRNIDmut1) and another with point mutations designed to disrupt its helical folding (NCoRNIDmut2) (Figure S9D-E). The interaction of these mutants with RR-DNA complexes was evaluated using MST experiments, showing no significant change in affinity for NCoRNIDmut1 compared to the wild type but a slight decrease for NCoRNIDmut2 (Table 1 and Figure S9F). These findings indicated that the IR region did not strongly contribute to the affinity between NCoRNID and RAR/RXR compared to CoRNR1 and CoRNR2. Altogether, our observations suggest that IR mainly interacts with RXR, in line with the minimal impact on affinity observed in RR-DNA complexes upon mutation. Furthermore, our HDX-MS data indicated that this interface is not responsive to the ligand or DNA, consistent with RXR’s repressive role, which remains primarily disconnected from RAR ligand or DNA interactions.

### Coactivator interaction dynamics with RAR/RXR and its DNA intercommunication

We next investigated whether DNA binding could also impact the dynamics of the RAR/RXR heterodimer in its active state, meaning in the presence of RAR and RXR agonist ligands. RAR/RXR heterodimer was prepared, bound to the different DNA binding sites in the presence of AM580 and CD3254, and subjected to HDX-MS (Figures S5, S6, and S8). By comparing the differential H/D exchange in RAR induced by DNA binding (Figure 4A), we observed predominant protection in the DBD, followed by mild, short-term protections in the hinge (Figure 4B). This result contrasts with the protections observed for apo-RAR (Figure 2J), likely reflecting differences in stabilization exerted by the ligands. The agonist ligand appeared to confer strong stabilization on RAR, leaving no room for additional stabilization by DNA (Figure S7D). Furthermore, the DR0 element remained the least stabilizing DNA under active conditions (Figure S7E), consistent with observations in apo-RAR. In addition, DNA binding induced significant protection in the RXR DBD and hinge region, strengthening the latter in the presence of the DR1 element (Figure 4C and S7E). Interestingly, DNA deprotection in the H2/H3 and H11 regions of the RXR LBD persisted in the active state. This deprotection, also observed in apo-RXR upon DNA binding, corresponds to the interface involved in coactivator interactions and could affect active complex formation.

**Figure 4.**
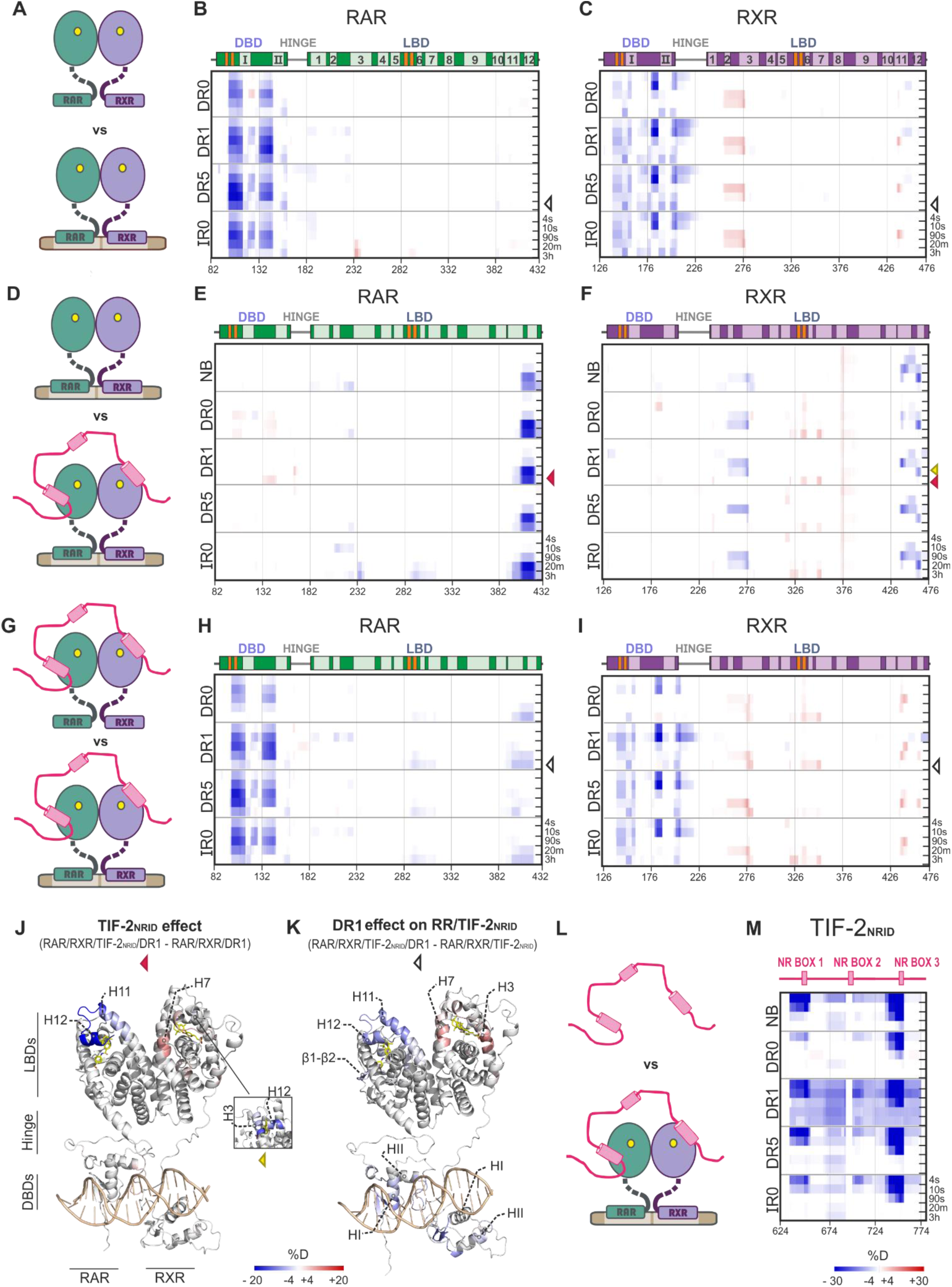
Differential HDX-MS reveals coactivator interaction on RAR and RXR and cooperativity with the DNA. **A)** Illustration of the active RAR/RXR vs active RAR/RXR bound to DNA. RAR is shown in green, while RXR is in purple. The yellow molecules represent the RAR (AM580) and the RXR (CD3254) agonist ligands. This color coding is used throughout the figures. **B)** Differential HDX-MS heatmap analysis on RAR for the agonist bound RR-DNAs (DR0, DR1, DR5 and IR0). Red shades indicate higher H/D exchange, suggesting regions of increased flexibility or exposure, whereas blue shades denote lower exchange, indicating structural protection. Secondary structure elements are shown above each heat map. The α-helices are represented in lighter colors followed by their respective numbering, while β-strands are shown in orange. **C)** Differential HDX-MS heatmap analysis of RXR for the agonist-bound RR-DNAs (DR0, DR1, DR5 and IR0). **D)** Illustration comparing the active RAR/RXR-DNA vs active RAR/RXR-DNA bound to TIF-2_NRID_. TIF-2_NRID_ is depicted in pink. The yellow molecules represent the RAR (AM580) and the RXR (CD3254) agonist ligands. **E)** Differential HDX-MS heatmap analysis of RAR, showing the impact of TIF-2_NRID_ interaction with RAR/RXR alone (NB) and in complex with the different DNAs. **F)** Differential HDX-MS data analysis of RXR, showing the impact of TIF-2_NRID_ interaction with RAR/RXR alone (NB) and in complex with different DNAs. **G)** Illustration of the active TIF-2_NRID_-RAR/RXR vs the active TIF-2_NRID_-RAR/RXR bound to the DNAs. **H)** Differential heatmap showing the changes in RAR upon DNA binding RAR/RXR/TIF-2_NRID_. **I)** Differential heatmap of the same conditions but for RXR. **J-K)** The HDX data plotted on the structure model generated based on the crystallographic structure of RAR/RXR-DR1 (5UAN pdb) (24). The data corresponds to exchange times of 3 hours and 20 minutes (small square) - highlighted on the right side of the heat maps by colored arrowheads. **J)** Active RAR/RXR-DR1 showing the differential HDX-MS effects caused by the presence of TIF-2_NRID_. **K)** RAR/RXR/TIF-2_NRID_ differential HDX-MS in the presence of the DR1 element. **L)** Illustration comparing TIF-2_NRID_ alone vs the TIF-2_NRID_ bound to RAR/RXR-DNA. **M)** Differential HDX-MS data analysis of TIF-2_NRID_, showing the impact of the binding of RAR/RXR unbound (NB) and bound to DNAs. Unexpectedly, RR-DR1 data appears noisy throughout the TIF-2_NRID_ sequence.

Subsequently, we evaluated how the variations in DNA binding could impact the coactivator interaction with RAR/RXR. We first measured the affinity between TIF-2NRID and the agonist-bound RR-DNA complexes by fluorescence anisotropy experiments (Table 1, Figure S2C). The dissociation constant (Kd) of the interaction between the RAR/RXR heterodimer and TIF-2NRID was in the nanomolar range, as previously measured for the RAR/RXR LBDs heterodimer (40). All the RR-DNA complexes had a significantly higher affinity for the coactivator than RR alone, suggesting that the DNA strengthens the RAR/RXR interaction with TIF-2NRID. This effect was more pronounced than for the interaction with NCoRNID. Interestingly, we also observed a lower affinity between TIF-2NRID and RR-DR0 compared to other RR-DNA complexes, which could be related to its transcriptional repressive role previously demonstrated in cellular contexts (7, 8).

To probe the interaction surfaces and the dynamic changes induced by the coactivator binding to RAR/RXR and to evaluate how DNA binding affects the coactivator interaction with the heterodimer, we completed the set of HDX-MS experiments on RAR/RXR and all the RR-DNA complexes prepared in the presence of TIF-2NRID (Figures S1E, 4D and G). We report the changes of H/D exchange in RAR/RXR and RR-DNA complexes owing to the presence of TIF-2NRID (Figure 4D-F) and in RR-TIF-2NRID owing to the presence of the different DNAs (Figure 4G-I). Our analysis showed that coactivator effects are limited to the LBD of both receptors. On RAR, adding TIF-2NRID to RR-DNA complexes generated strong protection of H12 (aa 392-417, Figure 4E and J). However, we did not observe any protection on H3 and H4, which, together with H12, are known to form a surface of interaction with the coactivator motif (11). This absence of protection is justified by the agonist ligand’s robust stabilization that primarily affected this region of RAR (Figure S7E), masking further HDX effects, differently from H12. Interestingly, the protection of H12 following TIF-2NRID addition was enhanced in all RR-DNA complexes compared to RAR/RXR alone (Figure 4E), suggesting that DNA plays a role in stabilizing the RR/TIF-2NRID complex. Adding DNA to the RR/TIF-2NRID complex also induced slight protection in the LBP region of RAR (β1/β2-H7 region) and more significantly in H11-H12, at late time points (Figure 4H and K). These observations suggest that DNA cooperats allosterically with the ligand for the coactivator binding to RAR/RXR. On RXR, the addition of TIF-2NRID caused protection on its H2-H3 (aa 243-273) and H11-H12 (aa 428-458) regions (Figure 4F), of much lower intensity than the ones observed for RAR H12, independently of the presence of DNA. These results demonstrated the coactivator interaction with both RAR and RXR, and the extent of protection highlighted RAR as the prominent binding site of TIF-2NRID in the heterodimer. Furthermore, we observed a deprotection in the β1/β2-H7 region of RXR following TIF-2NRID interaction, exclusively observed in RR-DNA complexes. The addition of DNA to the RR-TIF-2NRID complex also led to a deprotection of the RXR region of interaction with the coactivator (H3 and H11-H12) and the β1/β2-H7 region (aa 323-348) (Figure 4F and I). This result indicates that DNA destabilizes the ligand and coactivator interface on RXR and that, in the full complex, RXR has to adapt to its heterodimerization partner.

Next, we explored the effect of RAR/RXR and RR-DNA complexes on TIF-2NRID to highlight its interacting regions (Figure 4L). The strong protections to H/D exchange at short time points in NR boxes 1 and 3 and their flanking regions upon RAR/RXR addition suggested that TIF-2NRID physically interacted with the two agonist-bound receptors through these two NR boxes (Figure 4M). This represents the first structural observation of TIF-2NRID interacting with both agonist-bound receptors via its NR boxes 1 and 3, supporting a model of synergistic binding in which a single TIF-2NRID molecule spanned the liganded RXR and RAR within the heterodimer (68, 69). Moreover, the varying protection intensities observed at the heterodimer and coactivator interfaces suggested a scenario in which the TIF-2NRID NR box 3 strongly interacted with RAR LBD, while a weaker interaction occurred between RXR LBD and the coactivator NR box 1 (Figure 4E-F and M). We observed different protections of TIF-2NRID bound to the RR-DNA complexes. In particular, the TIF-2NRID interaction with RR-DR0 complex led to a significantly lower protection on both motifs (Figure 4M), agreeing with its lower affinity. In conclusion, we show that the binding of the heterodimer-DNA complexes to TIF-2NRID can be modulated by the DNA type and is in favor of a cooperative binding of NR boxes 1 and 3.

## Discussion

We combined several structural and biophysical methods with advanced molecular modeling to provide new insights into the dynamics and molecular interfaces at play in the RAR/RXR heterodimer, revealing correlations between conformational variations, DNA, ligand and coregulators binding. The excellent agreement between the SAXS intensity curves and the MD simulations provided us strong confidence in the analysis of the subtle differences between the different RARE-bound RAR/RXR heterodimers. Moreover, the use of ensembles to describe each of the studied conditions allowed us to capture the dynamics of individual domains within the RAR/RXR heterodimer. Rather than adopting exclusively compact or extended conformations, the RARE-bound RAR/RXR heterodimers presented an equilibrium among interdomain distances depending on the RARE sequences (Figure 5). Specifically, in DR0- and DR1-bound heterodimers, the RAR DBD and LBD preferentially adopted a close spatial arrangement, in agreement with the crystal structure of RARβ/RXR bound to DR1 (24) and the solution model for RARα/RXR bound to DR0 (39). Finally, although we identified transient contacts between the RAR LBD and DBD, aligned with the structural analysis of the DR1-bound complex, our study provides a more comprehensive picture of these complexes by describing the multiple states they can adopt due to their dynamic nature.

**Figure 5.**
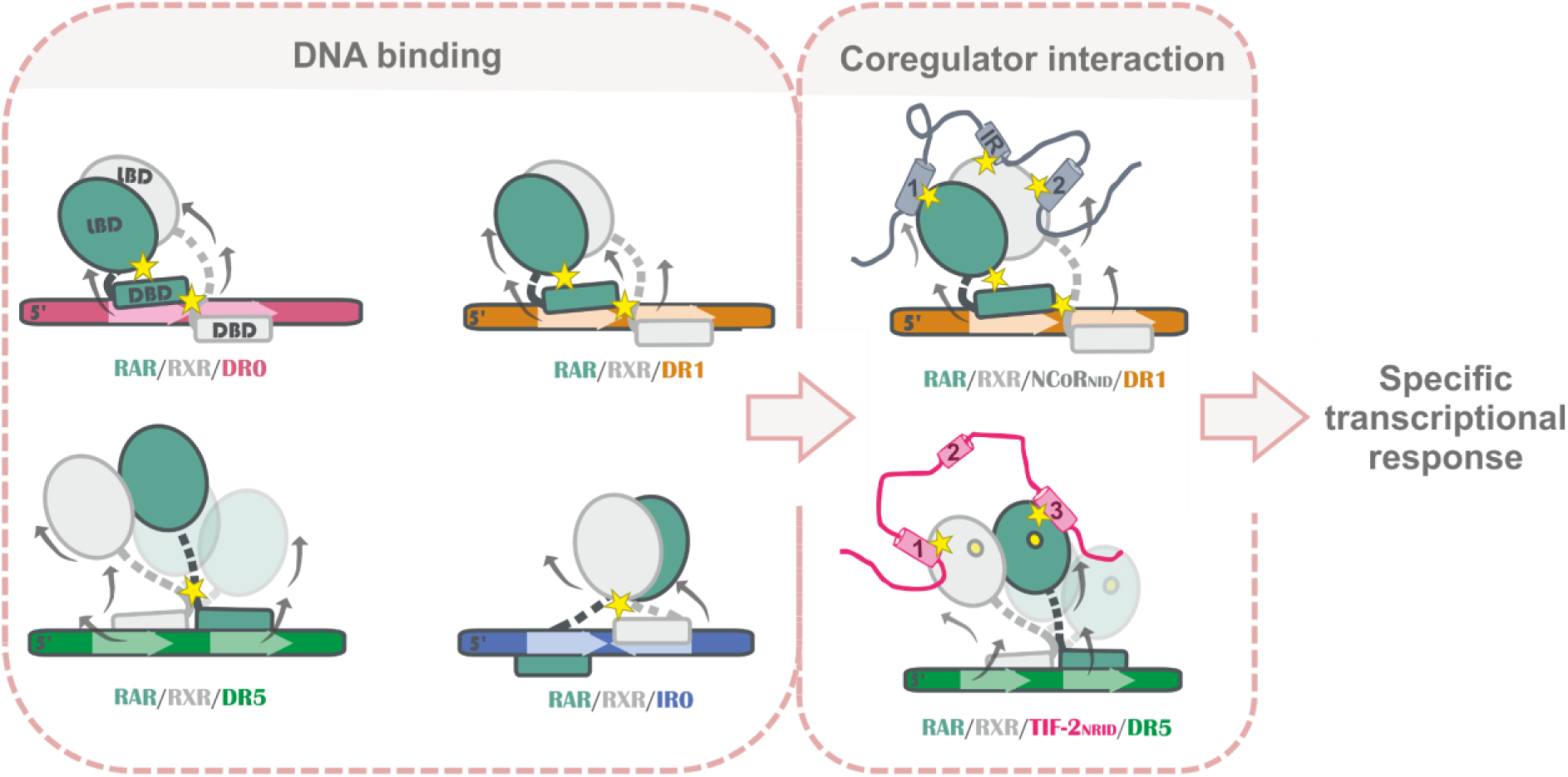
Interplay of DNA and Coregulator Binding in RAR/RXR Transcriptional Regulation. Schematic representation of RAR/RXR heterodimer DNA binding and conformational variations across different RAREs (DR0, DR1, DR5, and IR0). Stars indicate direct contacts, while arrows represent the allosteric effects induced by DNA binding across DBD and LBD domains. Coregulator interaction motifs are labeled numerically, corresponding to NCoR_NID_ (corepressor) and TIF-2_NRID_ (coactivator). Our results indicate that DNA binding drives conformational changes in RAR/RXR, establishing distinct patterns of interdomain contacts and allosteric stabilization across the heterodimer domains, assigning a unique pattern to each DNA-bound complex. The DNA-specific conformational signature is maintained upon coregulator binding, influencing its interaction dynamics and fine-tuning coregulator engagement, ultimately shaping the complex’s transcriptional response.

We also showed that DNA binding induced significant changes to solvent accessibility in the unliganded heterodimer LBDs, especially on the ligand binding pocket and in the region of interaction with coregulators. Furthermore, we observed that the RAR and RXR hinges established interdomain contacts depending on the DNA-bound complex. These observations reinforced the relevant role of hinges beyond merely providing flexibility, as recently demonstrated for full-length FXR (70), in transmitting information throughout the RAR/RXR complex. Previous studies on RARβ/RXR-DR1 and VDR/RXR have pointed out significant changes to HDX solvent accessibility in the hinge and the RAR or VDR DBDs induced by ligand binding, suggesting that ligand binding can orchestrate interdomain communication within RAR and VDR (24, 32). Here, we showed that not only ligand occupancy of the LBD, but more significantly, the RARE binding regulates interdomain communication between the RAR/RXR heterodimer domains.

In this work, we also provided the first structural evidence of DNA’s impact on coregulators complexed with nuclear receptors, despite the challenges posed by their highly dynamic and disordered nature. In the agonist bound complex, the stabilization of RAR/RXR mainly comes from the ligand, although the DNA provided an additional stabilization of RAR and a destabilization of RXR. A similar effect was observed by isothermal titration calorimetry (ITC) on affinity measurements with the RXR/PPARɣ heterodimer, where the DNA caused a reduction in the conformational entropy of the complex and energetically favored the interaction with the coactivator SRC-2NRID (31). Conversely, in in the unliganded complex, DNA played a significant role in stabilizing the heterodimer in interaction with the corepressor, with DR1 showing the strongest effect. This illustrats the allosteric translation of an event occurring in one domain (DBD) to another (LBD) towards an interactor (NCoRNID). Furthermore, a new region on NCoR, the IR, was revealed to interact with RXR. However, the precise role of this IR region remained unclear and warrants further investigation. In particular, we hypothesize that this IR may participate in additional molecular interactions within a larger complex, though its exact function in the cellular context remains unresolved. Finally, our results allowed us to designate RAR as the primary anchor point for coregulator interaction and the main sensor of the stabilization effects of ligands, DNA and coregulator within the heterodimer. On the contrary, RXR plays a subordinate role, emphasized in the presence of DNA, in agreement with its role as heterodimerization partner of many different NRs. Overall, our results provide a complete description of the structural dynamics in the transcriptional response regulated by RAR/RXR interaction with coregulators, detailing all the DNA direct and allosteric effects on it.

Extending structural studies on intact complexes comprising NR heterodimers, ligands and coregulator proteins could further advance our understanding of signal propagation within these functional assemblies and step further on their functioning. Whereas structural techniques such as X-ray crystallography and cryo-electron microscopy can be pivotal in deciphering the structures of NRs, integrative studies such as ours, combining molecular dynamics with solution methods, prove very powerful in describing ligand, DNA and coregulator-induced conformations and dynamics for these highly flexible NR complexes, and can be applied to enhance the understanding of other nuclear receptors beyond RAR/RXR.

## DATA AVAILABILITY

The structure has been deposited at the Protein Data Bank under the accession code (to be added). The mass spectrometry proteomics data have been deposited to the ProteomeXchange Consortium via the PRIDE (71) partner repository with the dataset identifier PXD061924. The SAXS data has been deposited at the SASDB under the accession code (to be added). Scripts used for MD simulation analysis are available in the GitHub repository (to be added).

## SUPPLEMENTARY DATA STATEMENT

Supplementary data are available at NAR online.

## ACKNOWLEDGEMENTS

We thank the SWING beamline at the SOLEIL synchrotron (Saint-Aubin, France) for regular beamtime allocation (Block Allocation Group, proposal No. 20201085) and assistance during data collection. We thank P. Germain for critical reading and discussion. We thank N. Sibille for the TIF-2NRID plasmid. We thank P. Poussin-Courmontagne for technical help, A. McEwen for help with X-ray data and the staff of Proxima 1 at synchrotron SOLEIL for assistance in using the beamline. We thank Christine Ueda from D.E. Shaw Research for providing the DES amber 3.20 force field files. We thank L. F. Rodrigues for generously providing the SAXS data plotting scripts and for their support in data analysis.

## AUTHOR CONTRIBUTIONS STATEMENT

ILT performed all experiments and data analysis and wrote the manuscript. AS conducted and analysed MD simulations. PV, DL, and PM supported HDX mass spectrometry experiments and data analysis. CC performed N-CoR_NID_ purification. NR carried out crystallography experiments and data analysis. WB contributed to critical reading and project conceptualization. PB provided support for SAXS data analysis and project conceptualization. ALM led project design and conceptualization, handled experiments, oversaw data analysis, and wrote the manuscript. All authors contributed to reviewing and editing the manuscript.

