## Supplementary material for "New structural insights into the control of the retinoic acid receptors RAR/RXR by DNA, ligands and transcriptional coregulators"

#### RAR/RXR DBDs purification, crystallization and structure resolution of RXR/RXR DBDs in complex with IR0

The HsRXRA DBD (130-212) and HsRARA DBD (82-167) were expressed in fusion with thioredoxine and hexahistidine tags. The proteins were produced and purified as described in (39). Fusion tags were removed by thrombin proteolysis and the cleaved proteins were then purified by SEC on a Superdex 75 in 10 mM HEPES pH 8, 50 mM NaCl, 5 mM $MgCl_2$, 1 mM TCEP. The HsRXR DBD, HsRAR DBD and DNA were mixed in an equimolar ratio and concentrated to a final concentration of 7 mg.mL$^{-1}$. The crystallization experiments were carried out by hanging drop vapor diffusion at 290 K by mixing equal volume (0.15 µl) of protein-DNA complex and of reservoir solution (20% PEG 1000, 0.2M MgCl2 0.1M NaCl, 0.05M Na cacodylate pH6.5). The crystals of the complex were transferred to artificial mother liquor containing 15 % Glycerol and flash cooled in liquid nitrogen. Data were collected on the Proxima 1 beamline of the synchrotron SOLEIL. The raw data were processed with XDS (71) and scaled with AIMLESS (72) programs. The structures were solved and refined using Phenix (73) and iterative model building using COOT (74). Data collection and refinement statistics are given in Table S4.

### SUPPLEMENTARY TABLES

**Supplementary Table 1:** List of DNA sequences used in this study.

| Element | Gene | Sequence | DNA length (base pair) |
|---|---|---|---|
| **DR0** | Hoxb13 | 5'-gaAGGTCAAGGCCAag-3' / 3'-ctTCCAGTTCCGGTtc-5' | 16 |
| **DR1** | Idealized | 5'-ctAGGTCAaAGGTCAgc-3'/ 3'-gcTGACCTtTGACCTag-5' | 17 |
| **DR5** | Rarb2 | 5'-agGGTTCAccgaaAGTTCAct-3'/ 3'-tcCCAAGTggcttTCAAGTga-5' | 21 |
| **IR0** | Trim16 | 5'- gcaGGGTCATGACCCcgc-3'/ 3'-gtCCCAGTACTGGGgcgc-5' | 18 |
| **DR0L** | Hoxb13 | 5'-cttccaggaAGGTCAAGGCCAagttgaaag-3'/ 3'-ctttcaactTGGCCTTGACCTtcctggaag-5 | 30 |

**Table S2.** Estimated parameters from SAXS intensities measured on RAR/RXR heterodimer bound to the different DNA binding sites (DR0, DR0L, DR1, DR5, IR0) in the presence and absence of NCORNID.

| Sample | Rg (nm) | | Estimated MM (kDa) | | Dmax (nm) | MMseq (kDa) | Ensemble fit to data (χ2) |
|---|---|---|---|---|---|---|---|
| | Guinier | p(r) function | Bayesian | Credibility Interval | | | |
| RR-DR0 | 3.6± 0.03 | 3.7± 0.02 | 78 | 73-84 | 12.7 | 87 | 1.0 |
| RR-DR0L | 4.1 ± 0.02 | 4.2± 0.1 | 79 | 73-84 | 13.9 | 95 | 1.4 |
| RR-DR1 | 3.8 ± 0.00 | 3.9 ± 0.06 | 86 | 84-95 | 12.4 | 88 | 1.4 |
| RR-DR5 | 3.9 ± 0.04 | 4.0 ± 0.02 | 86 | 84-95 | 12.7 | 90 | 1.0 |
| RR-IR0 | 3.9 ± 0.01 | 3.9 ± 0.07 | 83 | 75-86 | 12.6 | 78 | 1.3 |
| NRR-DR0L | 5.4 ± 0.03 | 5.5± 0.4 | 208 | 176-221 | 20.1 | 126 | 1.6 |
| NRR-DR1 | 5.6 ± 0.03 | 6.0 ± 0.3 | 170 | 151-176 | 21.7 | 118 | 2.1 |
| NRR-DR5 | 5.7 ± 0.06 | 6.0 ± 0.4 | 186 | 162-194 | 21.7 | 120 | 1.5 |
| NRR-IR0 | 5.5 ± 0.04 | 6.0 ± 0.5 | 186 | 162-194 | 22.9 | 108 | 1.3 |

**Table S3.** Time-resolved fluorescence parameters of the RAR-RXR-OG488 in the absence and in the presence of saturating concentrations of the different DNA-TAMRA5/6. FRET efficiency at saturation (E) was calculated as described in (39).

| RAR/RXR | E |
|---|---|
| RARb2 DR5* | 0.46 |
| Ramp2 DR1* | 0.75 |
| Hoxb13 DR0* | 0.81 |
| Trim16 IR0 | 0.69 |

**Table S4.** X-ray crystallography data collection and refinement statistics.

| | **RXR DBD - IR0** |
|---|---|
| **Data collection** | |
| Beamline | PX1 |
| Space group | P $4_1$ $2_1$ 2 |
| Unit-cell parameters (Å, °) | 56.019 56.019 169.871, 90.00 90.00 90.00 |
| Resolution range (Å) | 33.84-3.5 |
| Unique reflections | 3712 |
| $CC_{1/2}$ | 1 |
| Completeness | 96.50 |
| **Refinement** | |
| Resolution range (Å) | 33.84-3.5 |
| $R_{work}$ / $R_{free}$ (%) | 26.53 / 33.5 |
| Number of non-hydrogen atoms | |
| macromolecules | |
| ligands | 1684 |
| | 4 |

### SUPPLEMENTARY FIGURES

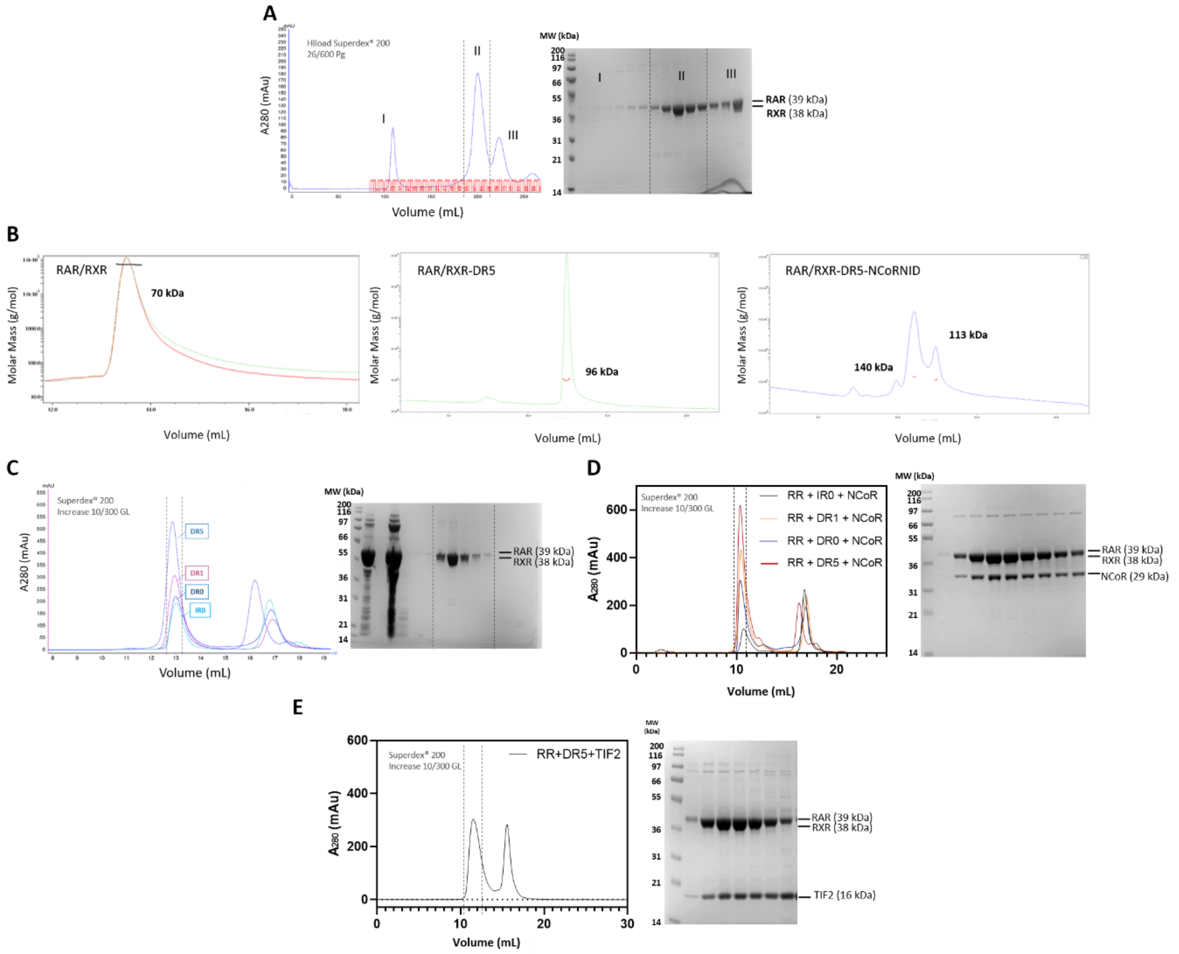


**Supplementary Figure 1. Sample preparation steps.** A) SEC chromatogram of RAR/RXR followed by the 10% SDS-page gel containing the representative fraction from some aggregated sample (peak I), the heterodimer in the stoichiometry of 1:1 (peak II) followed by an excess of RAR monomer (peak III). B) SEC-MALS chromatogram showing the scattered light profile for the heterodimer alone (RAR/RXR, left), the heterodimer bound to DR5 (RAR/RXR-DR5, middle) and the repressive complex (RAR/RXR-DR5-NCoR$_{NID}$, right). C) SEC chromatogram of RAR/RXR bound to the different DNAs. The complex eluted in the first peak (dotted line), and the second peak refers to the excess of DNA added during previous incubation. The variation in the elution volume reflects the slight difference in the DNA length. The sharp peak in the expected elution volume (13 mL - by using a Superdex 200 Increase 10/300) together with the 10% SDS-page gel confirms the homogeneity and purity of the sample. D) SEC chromatography RAR/RXR bound to four different DNAs (DR0, DR1, DR5 and IR0) and the corepressor NCoR$_{NID}$ - which, despite its lower molecular weight, it is migrating above the 35 kDa marker. The elution volume (~11 mL) reveals the shift provoked by corepressor binding. The SDS-PAGE gel confirms the complex formation and its purity. E) SEC chromatography RAR/RXR bound to the DR5 element and the coactivator TIF-2$_{NRID}$. The elution volume (~11.5 mL) reveals the shift caused by the complex formation with the coactivator. The SDS-PAGE gel confirms the complex formation and its purity.

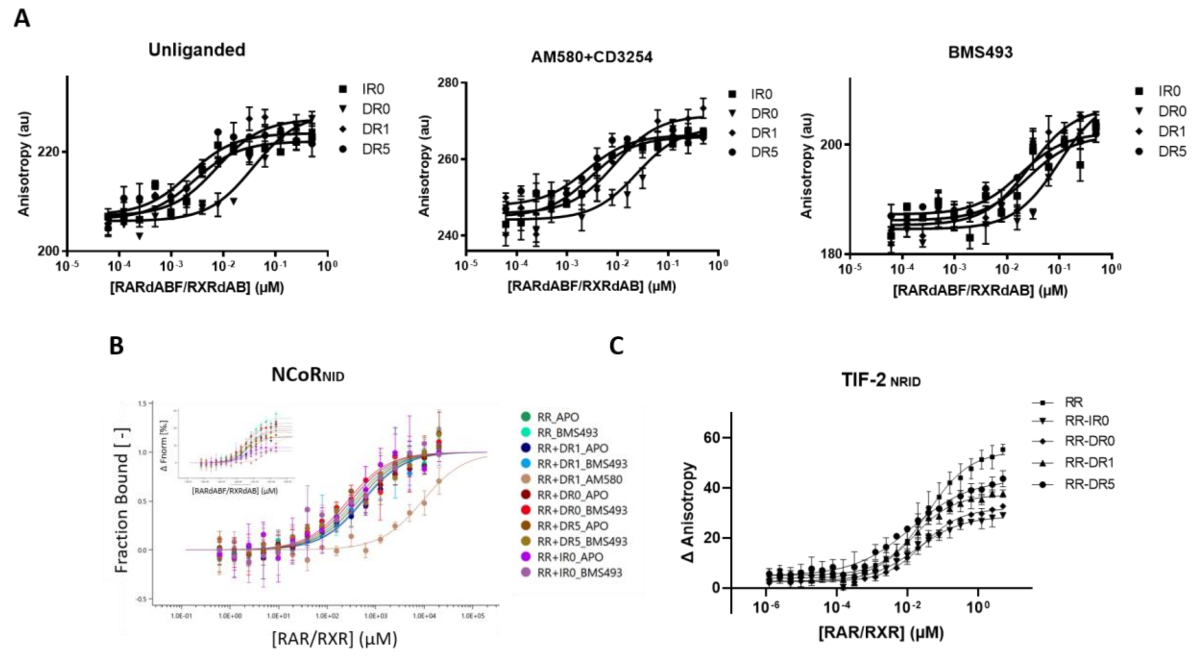


**Supplementary Figure 2. Titration curves of the affinity measurements on RAR/RXR heterodimer.** A) Affinity curves obtained by fluorescence anisotropy for heterodimer RAR/RXR and the different DNAs (IR0, DR0, DR1 and DR5) in the absence of the ligand (left), or in the presence of the agonists AM580 and CD3254 (middle), or the presence of the inverse agonist BMS493 (right). B) Affinity curves measured by microscale thermophoresis of the RAR/RXR unbound (RR) and bound to the different DNAs (RR-IR0, RR-DR0, RR-DR1, RR-DR5) with the corepressor NCoR$_{NID}$, in the presence and absence (apo) of the inverse agonist BMS493. C) Affinity curves by fluorescence anisotropy for the RAR/RXR unbound (RR) and bound to the different DNAs (RR-IR0, RR-DR0, RR-DR1, RR-DR5) with the coactivator TIF-2$_{NRID}$. All the measurements were performed in the presence of the RARα and RXR respective agonists, AM580 and CD3254.

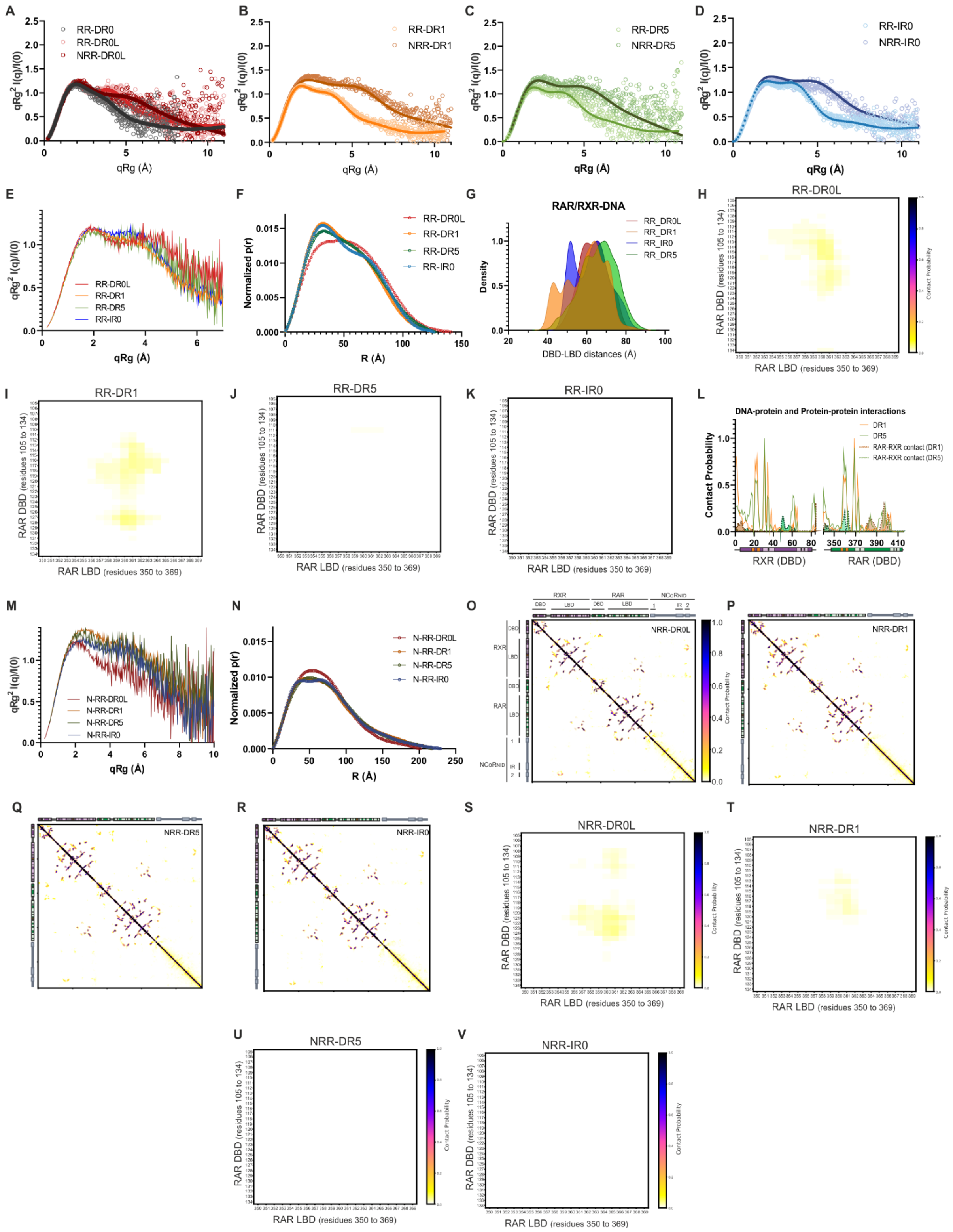


**Supplementary Figure 3. SAXS and molecular dynamics analysis of RAR-RXR complexes bound to different DNA binding sites in the presence and absence of $NCoR_{NID}$.** (A-D) Kratky plots derived from SAXS for RAR/RXR complexes bound to the different DNA binding sites in the absence (RR) and presence of $NCoR_{NID}$ (NRR). Panel (A) shows RR-DR0 (grey), RR-DR0L (red), and N-RR-DR0L (dark red). Panel (B) depicts RR-DR1 (orange) and N-RR-DR1 (dark orange); panel (C) displays RR-DR5 (green) and N-RR-DR5 (dark green); and panel (D) presents RR-IR0 (blue) and N-RR-DR0 (dark blue). (E) Superimposed Kratky plots for RR complexes bound to the distinct DNA binding sites (DR0L, DR1, DR5, IR0). (F) Superposition of pairwise distance distribution functions P(r) for RR-DNA complexes. (G) Superposition of distances calculated between the centers of mass of the DBD and LBD for RR-DNA complexes. (H-K) Contact probability submatrix for the RAR DBD and LBD regions (ranging from 105-134 and 350-369 residues, respectively) showing the transient interdomain contacts on RAR bound to DR0L and DR1 predominantly. Panels correspond to the respective DNA binding sites: (H) DR0L; (I) DR1; (J) DR5; (K) IR0. (L) Contacts between the DBDs of RAR and RXR with DNA in RR complexes bound to DR1 (yellow) and DR5 (green). The dotted lines indicate inter-subunit DBD interactions: dark yellow for DR1 and green for DR5. (M) Superimposed Kratky plots for N-RR-DNA complexes. (N) Superposition of P(r) plots obtained for N-RR-DNA complexes. (O-R) Contact probability maps for NCoRNID-RAR/RXR complexes from the molecular dynamics simulations after re-weighting process. Proteins' secondary structures are indicated by colored bars: RXR (purple) and RAR (green), and NCoRNID (lilac), with alpha-helices (light shades) and beta-sheets (orange). Panels correspond to the DNA binding sites: DR0L (O) DR1 (P), DR5 (Q), and R0 (R). (S-V) Contact probability submatrix for the RAR DBD and LBD regions (ranging from 105-134 and 350-369 residues, respectively) within the RAR/RXR/NCoR/DNAs complexes showing the transient interdomain contacts on RAR bound to DR0L and DR1 predominantly. Panels correspond to the respective DNA binding sites: (S) DR0L; (T) DR1; (U) DR5; (V) IR0.

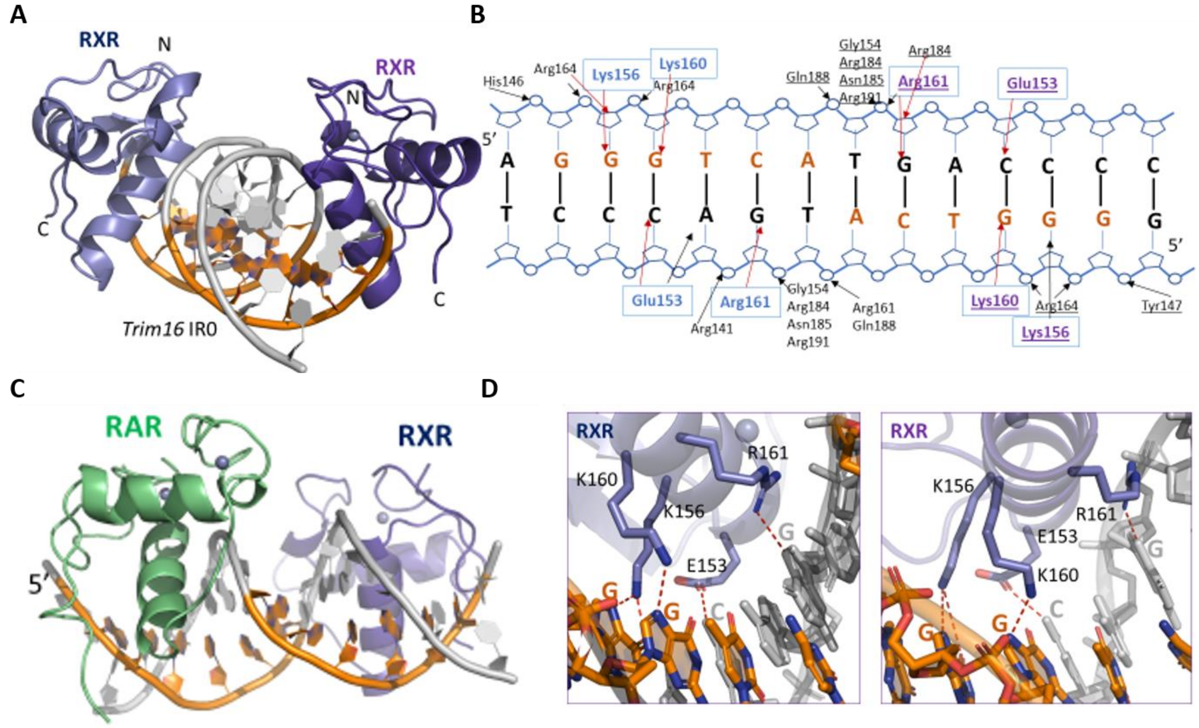


**Supplementary Figure 4.** Crystal structure of RXR–RXR DBDs-Trim16 IR0. (A) Overall structure of RXR–RXR–DNA complex. The spheres indicate the Zn atoms. (B) Schematic view of the RXR–RXR DBDs-Trim16 IR0 contacts calculated with NUCPLOT with a 3.9 Å distance cutoff. (C) Specific interactions of RXR homodimer DBDs to Trim16 IR0. Left: View along the DNA-recognition helix of 5' RXR. Right: The corresponding view of 3’ RXR. Hydrogen-bonds are shown as red dotted lines. (D) 3D model of RAR-RXR DBDs bound to Trim16 IR0 with RAR bound to the 5’ half site based on time-resolved fluorescence parameters.

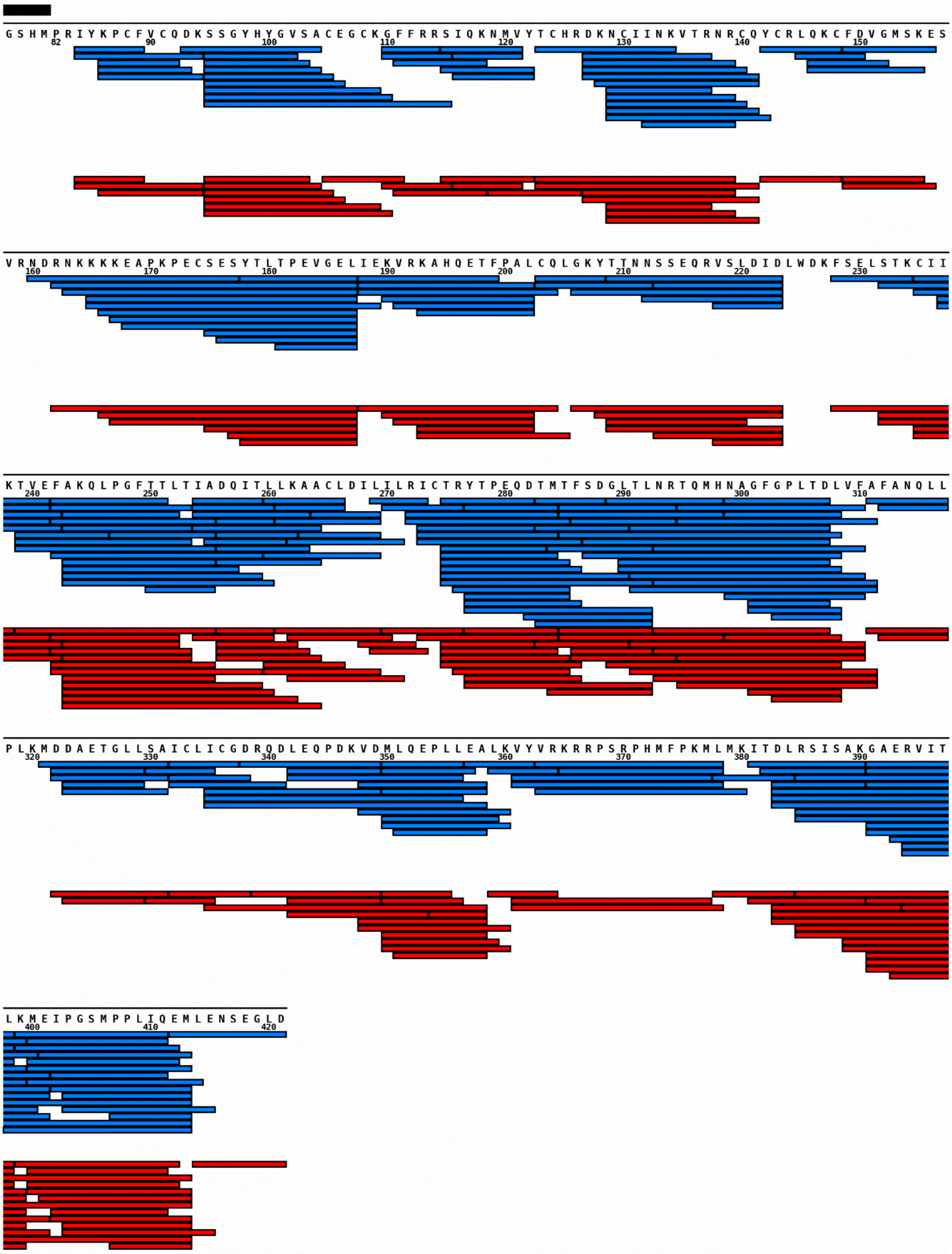


**Supplementary Figure 5. Sequence coverage of RAR.** Coverage map of RAR in the experiment with TIF (blue bars) and NCoR (red bars). Digestion parameters for TIF-2/NCoR data were - sequence coverage: 93.4%/92.6%. Number of peptides: 210/147. Average peptide length: 11.7/11.7. Average redundancy: 7.5/5.3. Only peptides providing HDX data are shown. The black box highlights the sequence from the tag. The map was plotted using MSTools (https://peterslab.org/MSTools/DrawMap/DrawMap.php) (56).

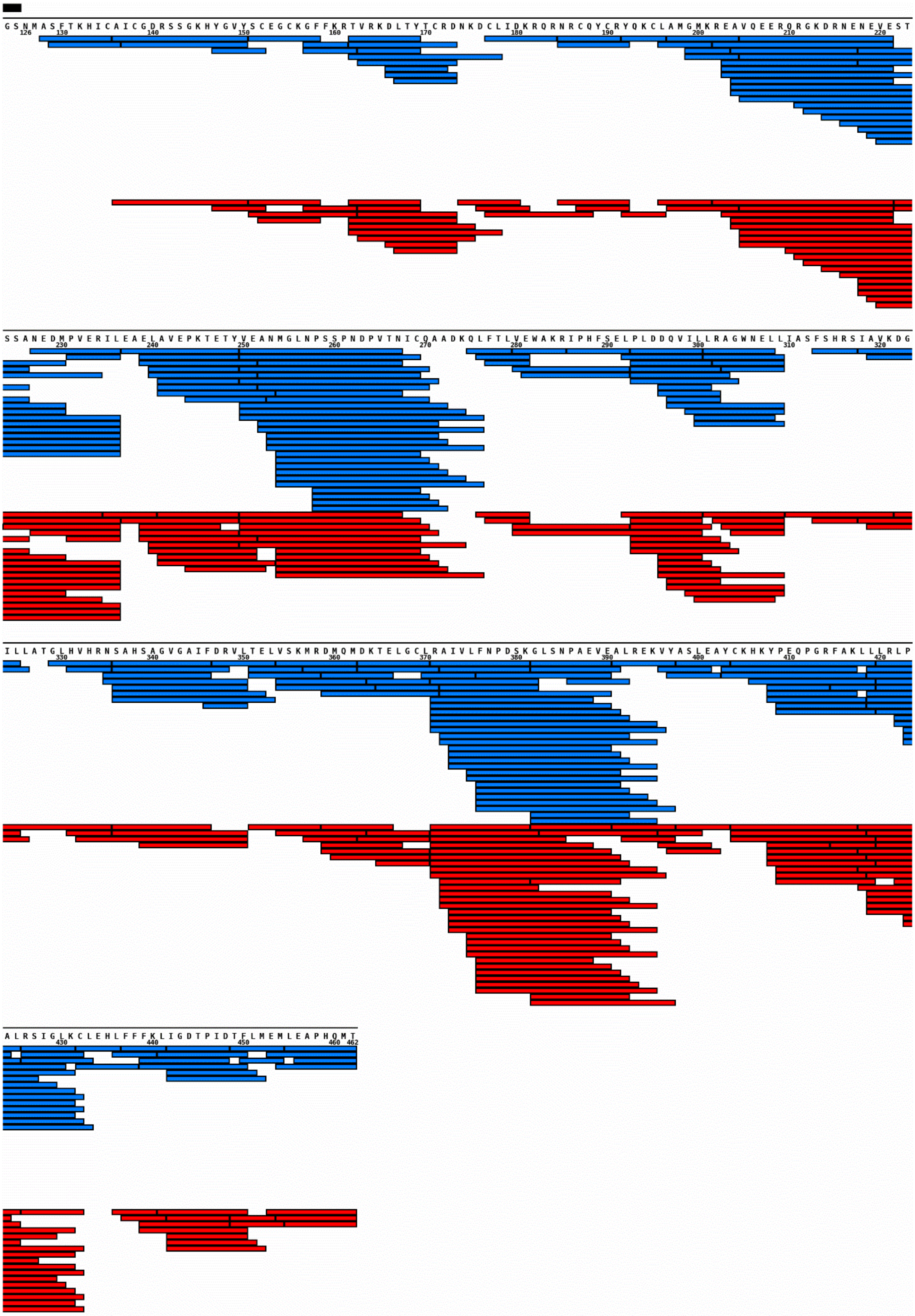

**Supplementary Figure 6. – Sequence coverage of RXR.** Coverage map of RXR in the experiment with TIF (blue bars) and NCoR (red bars). Digestion parameters for TIF-2/NCoR data were - sequence coverage: 95.4%/93.6%. Number of peptides: 210/192. Average peptide length: 12.6/12.5. Average redundancy: 8.0/7.4. Only peptides providing HDX data are shown. The black box highlights the sequence from the tag. The map was plotted using MSTools (https://peterslab.org/MSTools/DrawMap/DrawMap.php) (56).

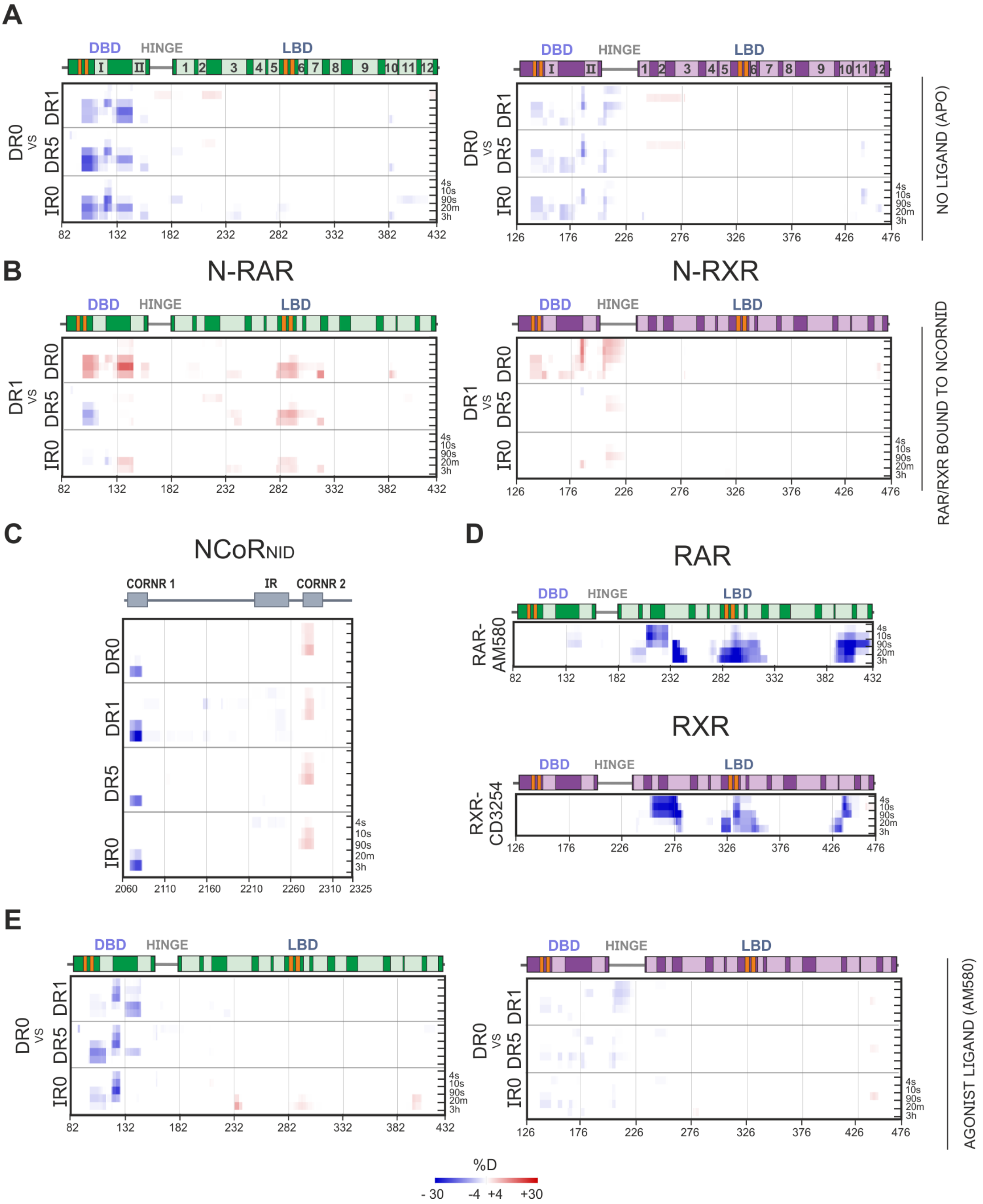


**Supplementary Figure 7. Further details on the differential protection on RAR/RXR heterodimer and NCoRNID caused by the DNA and ligands.** (A) Heatmap of the apo RAR (left) and RXR (right) showing the differential in protection between DR0 and the other DNA conditions, such as bound to DR1 (DR1 - DR0), DR5 (DR5 - DR0) and IR0 (IR0 - DR0). Red and blue shades represent increased and decreased deuterium exchange, respectively. The α-helices are represented in lighter-colored rectangles followed by their respective numbering. β-strands are represented in orange. (B) Heatmap of the RAR (left) bound to the inverse agonist and corepressor (left) and apo RXR in the presence of the corepressor (right) showing the differential in deuterium incorporation caused by the DR1 binding compared among the other DNA binding sites. (C) Differential HDX-MS heatmap analysis on NCoR$_{NID}$ showing the DNA-ligand effect on increasing the protection on its CoRNR1 and decreased on the CoRNR2. (D) Heatmaps showing the protection caused by the agonist ligand AM580 interaction with the RAR (upper) and RXR (lower). (E) Differential heatmap of the agonist bound RAR (AM580, left) and RXR (CD3254, right) showing the variation in protection between DR0 and the other DNA conditions, such as bound to DR1 (DR1 - DR0), DR5 (DR5 - DR0) and IR0 (IR0 - DR0).

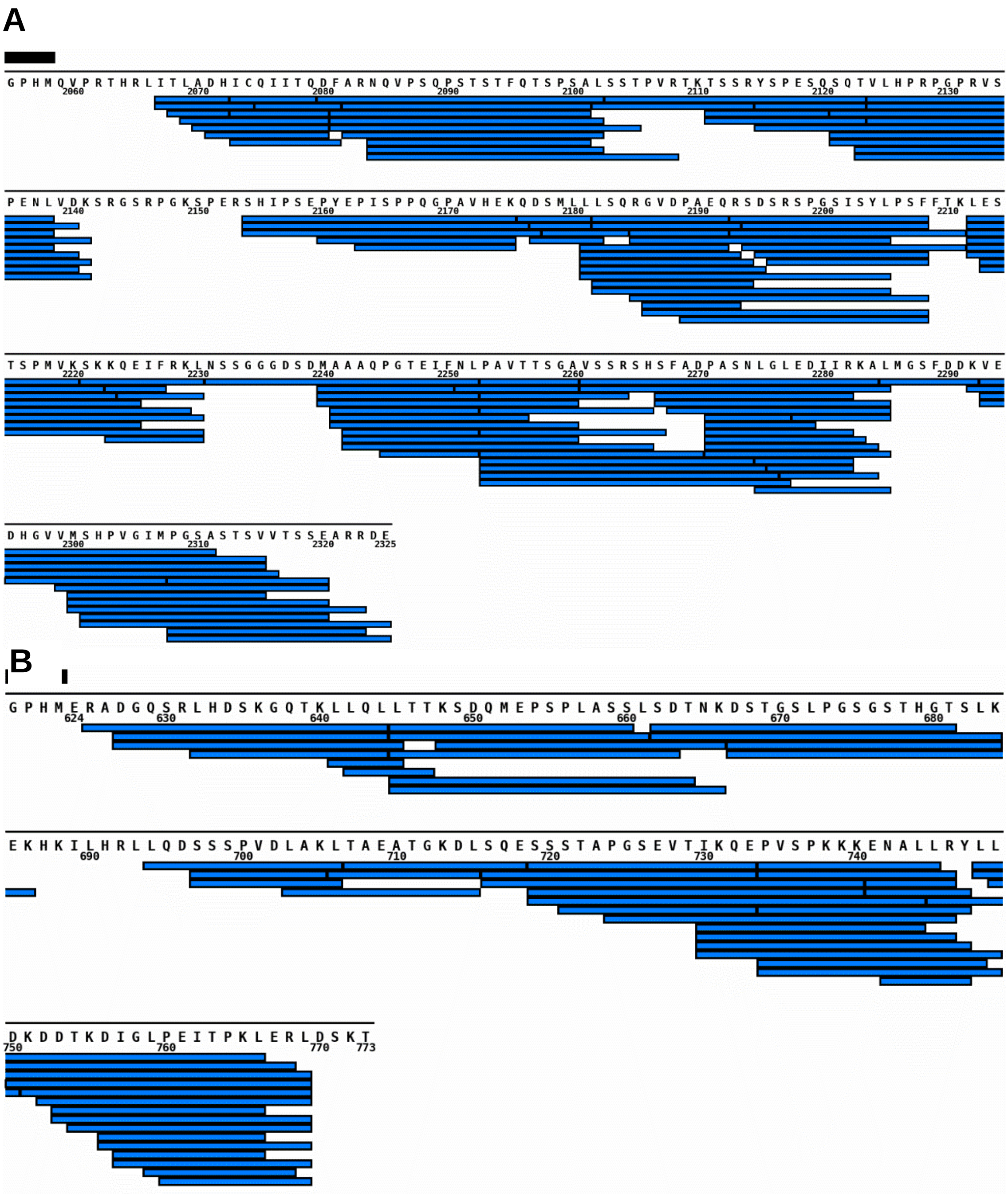

**Suppleementary Figure 8. Sequence coverage of NCoR and TIF2.** Coverage map of NCoR (A) and TIF-2(B). Digestion parameters for NCoR were - sequence coverage: 89.54%, number of peptides: 127, average peptide length: 15.5, average redundancy: 8.0. Digestion parameters for TIF-2 were - sequence coverage: 87.8%, number of peptides: 57, average peptide length: 15.5, average redundancy: 6.4. Only peptides providing HDX data are shown. The black box highlights the sequence from the tag. The map was plotted using MSTools (https://peterslab.org/MSTools/DrawMap/DrawMap.php) (56).

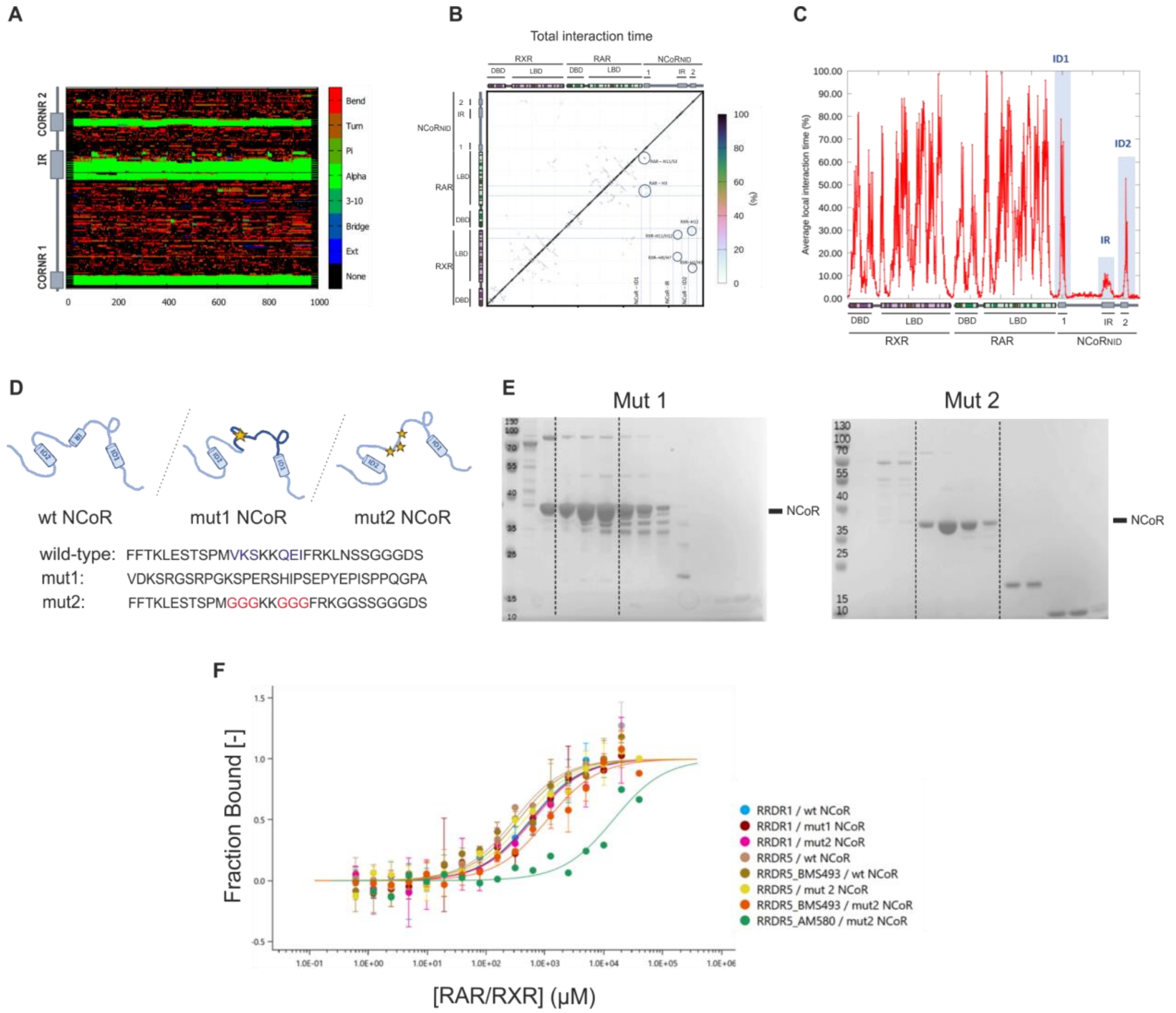

**Supplementary Figure 9. NCoR$_{NID}$ mutants sample preparation and affinities with the RAR/RXR-DNA complexes.** A) Secondary structure proportion of the RAR/RXR/NCoRNID-DR1 (RRN-DR1) complex through molecular simulation. B) Contact map among the proteins in the RRN-DR1 complex. C) Average local interaction time (%) within the RRN-DR1. D) Illustration of the two mutations designed for NCoR$_{NID}$ on its intermediate region (IR). NCoR$_{NID}$mut1 is composed of a complete replacement of the IR on the NCoR$_{NID}$. NCoR$_{NID}$mut2 is composed of punctual mutations to glycines on crucial points for IR helicity, potentially causing the disruption of its secondary structure as well as its motif of interaction with the heterodimer. E) SDS-PAGE after the last step of purification done by SEC for the NCoR$_{NID}$mut1 (left) and NCoR$_{NID}$mut2 (right), showing the suitable sample purity for both preparations. F) Affinity curves measured by microscale thermophoresis of the RR-DNA complexes with the corepressor NCoR$_{NID}$mut1 and 2, in the presence and absence (apo) of the inverse agonist BMS493 or the RARα agonist (AM580).
